# A visual encoding model links magnetoencephalography signals to neural synchrony in human cortex

**DOI:** 10.1101/2020.04.19.049197

**Authors:** Eline R. Kupers, Noah C. Benson, Jonathan Winawer

## Abstract

Synchronization of neuronal responses over large distances is hypothesized to be important for many cortical functions. However, no straightforward methods exist to estimate synchrony non-invasively in the living human brain. MEG and EEG measure the whole brain, but the sensors pool over large, overlapping cortical regions, obscuring the underlying neural synchrony. Here, we developed a model from stimulus to cortex to MEG sensors to disentangle neural synchrony from spatial pooling of the instrument. We find that synchrony across cortex has a surprisingly large and systematic effect on predicted MEG spatial topography. We then conducted visual MEG experiments and separated responses into stimulus-locked and broadband components. The stimulus-locked topography was similar to model predictions assuming synchronous neural sources, whereas the broadband topography was similar to model predictions assuming asynchronous sources. We infer that visual stimulation elicits two distinct types of neural responses, one highly synchronous and one largely asynchronous across cortex.

## 1. Introduction

Two of the most widely used tools to study dynamic cognitive processes in the human brain are magnetoencephalography (MEG) and electroencephalography (EEG). These measurement techniques provide whole brain coverage at millisecond time resolution, allowing researchers to extract complex spatiotemporal dynamics non-invasively in both healthy and clinical populations. The electric and magnetic fields measured by these instruments reflect the superposition of all cellular processes that generate current. These processes span temporal scales from a millisecond or less (*e.g.*, action potentials) to slow cortical potentials (< 1 Hz; (Birbaumer, Elbert, Canavan, & Rockstroh, 1990)); they span cellular scales from dendritic spines to axons and dendrites; and they span circuitry scales, from the idiosyncratic fluctuations in the membrane potential of a single neuron to highly synchronized responses across centimeters of brain tissue (Buzsaki, Anastassiou, & Koch, 2012).

Much of the MEG and EEG literature has focused on neural responses that are synchronized across extended regions of cortex. Widespread neural synchrony has been observed in many tasks and states. For example, rhythmic responses in the field potential, believed to reflect underlying widespread synchrony, often accompany sensory information processing (Buzsaki et al., 2012). Changes in cortical states are often characterized by changes in synchrony (such as alertness versus asleep (Steriade, McCormick, & Sejnowski, 1993)), and neurological disorders can be correlated with changes in neural synchrony (Uhlhaas & Singer, 2006).

It is thus important to characterize neural synchrony with non-invasive tools. In recent decades, there have been substantial advances in measurement methodology such as high-density EEG (Robinson et al., 2017) and on-scalp MEG sensor arrays (Iivanainen, Stenroos, & Parkkonen, 2017), and in biophysical forward modeling of neuronal currents to extracranial sensor responses (Stenroos & Nummenmaa, 2016). Nonetheless, it is still not possible to unambiguously infer the spatiotemporal pattern of neural source activity from the measured sensor responses. This is because each sensor pools over large and overlapping cortical regions, resulting in many possible combinations of source activity that could explain any particular observed pattern of sensor responses (Hämäläinen, Hari, Ilmoniemi, Knuutila, & Lounasmaa, 1993). In some cases, increases in power measured extracranially are explained by greater coherence across cortex (but, on average, *decreased* power in cortical fluctuations (Musall, von Pfostl, Rauch, Logothetis, & Whittingstall, 2014)). In other cases, increases in power measured extracranially are explained by increases in the power of local cortical fluctuations without accompanying widespread synchronization (Frauscher, von Ellenrieder, Dubeau, & Gotman, 2015).

One way to make inferences about the spatiotemporal pattern of neuronal sources giving rise to MEG or EEG data is to invert the biophysical forward model. Because there are many more neural sources than there are sensors, the problem is ill-posed (there are many solutions). Typically, researchers arrive at a single solution by applying a regularizer or other constraints. For example, one can choose the source activity solution with the smallest L2-norm (*e.g.* with minimum norm estimation (Hamalainen & Ilmoniemi, 1994)). This may be appropriate when there is no *a priori* knowledge of the likely pattern of source activity. However, the assumptions implicit in the regularizers are, at best, an approximation, and in some cases may be highly inaccurate. For example, regularizers will penalize “silent sources”, such as simultaneous neural responses from opposite-facing dipoles, even though such neural responses may be present and even the largest source of activity. In the extreme, seizures are defined by widespread and highly synchronous neural activity. Accurate interpretation of the cortical sources of seizure activity has important health implications and is an active area of study (Acar, Makeig, & Worrell, 2008).

Here, we present an alternative approach that makes use of prior knowledge to predict sensor responses from visual stimulation by combining an encoding model from stimulus to cortex with a forward model from cortex to sensors (**Figure 1**). This approach allows us to separate the contribution of neural synchrony on the cortex from the pooling function of the sensors. By doing so, we can test specific but constrained hypotheses about the spatiotemporal pattern of neuronal responses.

**Figure 1.**
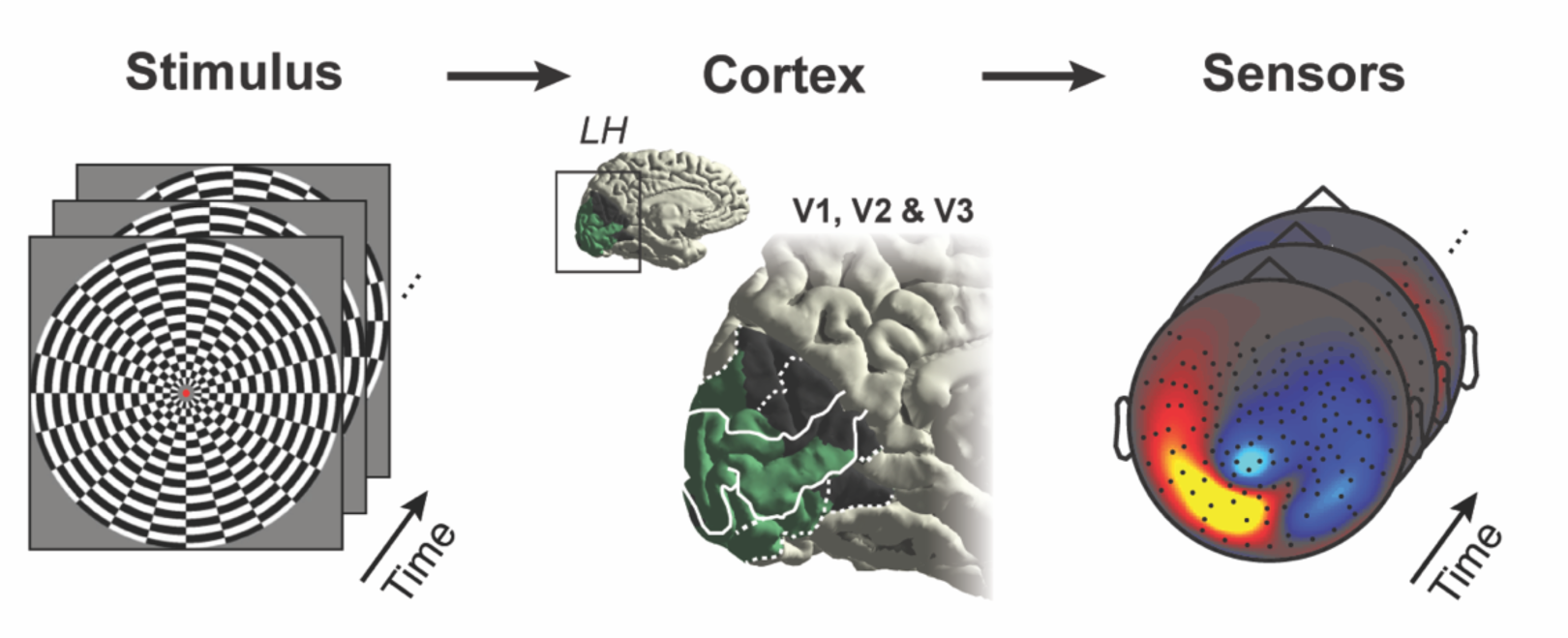
A visual encoding model for MEG. The modeling approach predicts sensor responses starting from a visual stimulus as input. The first stage predicts cortical responses from the stimulus. The second stage predicts sensor responses from the cortical activity. For illustration purposes, only the left hemisphere (LH) is depicted. (The forward model uses both hemispheres.)

Specifically, we examine two components of the MEG response to visual stimuli which are believed to reflect different kinds of neural processes with different degrees of cortical synchrony. One component is a *stimulus-locked response*, fluctuations in the MEG signal at the frequency of stimulus contrast-reversals. The second component is a *broadband response*, a spectrally broad increase in amplitude, including temporal frequencies that are not in the stimulus. The stimulus-locked response is often measured extracranially (reviewed by (Norcia, Appelbaum, Ales, Cottereau, & Rossion, 2015)), whereas broadband responses are more typically measured intracranially (Miller et al., 2014). There has been little study of the two signal components in the same individuals in the same experiments with noninvasive methods. Here, we measure both of these signal components in the same subjects and compare the spatial patterns of the responses across the sensor array to patterns predicted by model-based simulations. Comparing the data to simulations with known ground truth helps to make inferences about the processes that give rise to the data.

To generate the two types of responses, we conducted a visual steady-state MEG experiment where subjects viewed a large high contrast-reversing dartboard pattern. This is a widely used paradigm to study stimulus-locked responses, also known as the steady-state visually evoked field (‘SSVEF’) or steady-state visually evoked potentials in EEG (‘SSVEP’) (Adrian & Matthews, 1934; Van Der Tweel & Lunel, 1965; Norcia & Tyler, 1985; Regan, 1989; Norcia et al., 2015). The steady-state paradigm with simple, high-contrast patterns has also been shown to be effective for eliciting broadband responses, both in intracranial measurements (Winawer et al., 2013) and MEG measurements (Kupers et al., 2018). While present in the gamma band, broadband responses show different response patterns compared to narrow-band gamma oscillations, for example to visual grating stimuli (Henrie & Shapley, 2005; Ray & Maunsell, 2011; Hermes, Miller, Wandell, & Winawer, 2015; Bartoli et al., 2019). Increased broadband power has been characterized across the brain (Crone, Miglioretti, Gordon, & Lesser, 1998; Miller et al., 2007; Miller, Sorensen, Ojemann, & den Nijs, 2009; Miller et al., 2014), is thought to be generated by different neural circuits compared to evoked responses (Manning, Jacobs, Fried, & Kahana, 2009; Miller et al., 2009; Milstein, Mormann, Fried, & Koch, 2009), and correlates well with the fMRI BOLD signal (Mukamel et al., 2005; Hermes et al., 2012; Hermes, Nguyen, & Winawer, 2017). Both of these signal components are likely to be important for understanding how cortical circuits encode visual information.

## 2. Results

### 2.1 Visually driven MEG response can be separated into a stimulus-locked and broadband component

Subjects viewed a large-field (22° diameter) dartboard pattern that contrast-reversed 12 times per second, interspersed with blanks (zero-contrast, mean luminance). We separated the sensor responses into two components, one time-locked to the stimulus (*stimulus-locked*) and one that is not (*broadband*), using the same method as in (Kupers et al., 2018). Because the acquisition and analysis methods were the same as in the prior study, we combined data from that study (N=6) with the newly acquired data (N=6), for a total of 12 datasets.

The stimulus-locked response tends to be large in visually responsive sensors and is defined as the difference in amplitude between stimulus and blank periods at 12 Hz (**Figure 2**, left panel). The second component is a spectrally broad increase in amplitude (**Figure 2**, right panel). The broadband response is defined as the elevation in amplitude over baseline from 60 to 150 Hz (excluding harmonics of the contrast-reversal rate). By definition, the broadband signal is <u>not</u> time-locked to the contrast-reversal rate of the stimulus (12 Hz or its harmonics). This method of separating the responses differs from the more conventional method of summing the amplitude (or power) within distinct temporal frequency bands. For example, 83 Hz and 84 Hz would both be considered part of the gamma band, but we include 83 Hz and exclude 84 Hz in the broadband computation, since 84 Hz is a harmonic of the contrast-reversal rate.

**Figure 2.**
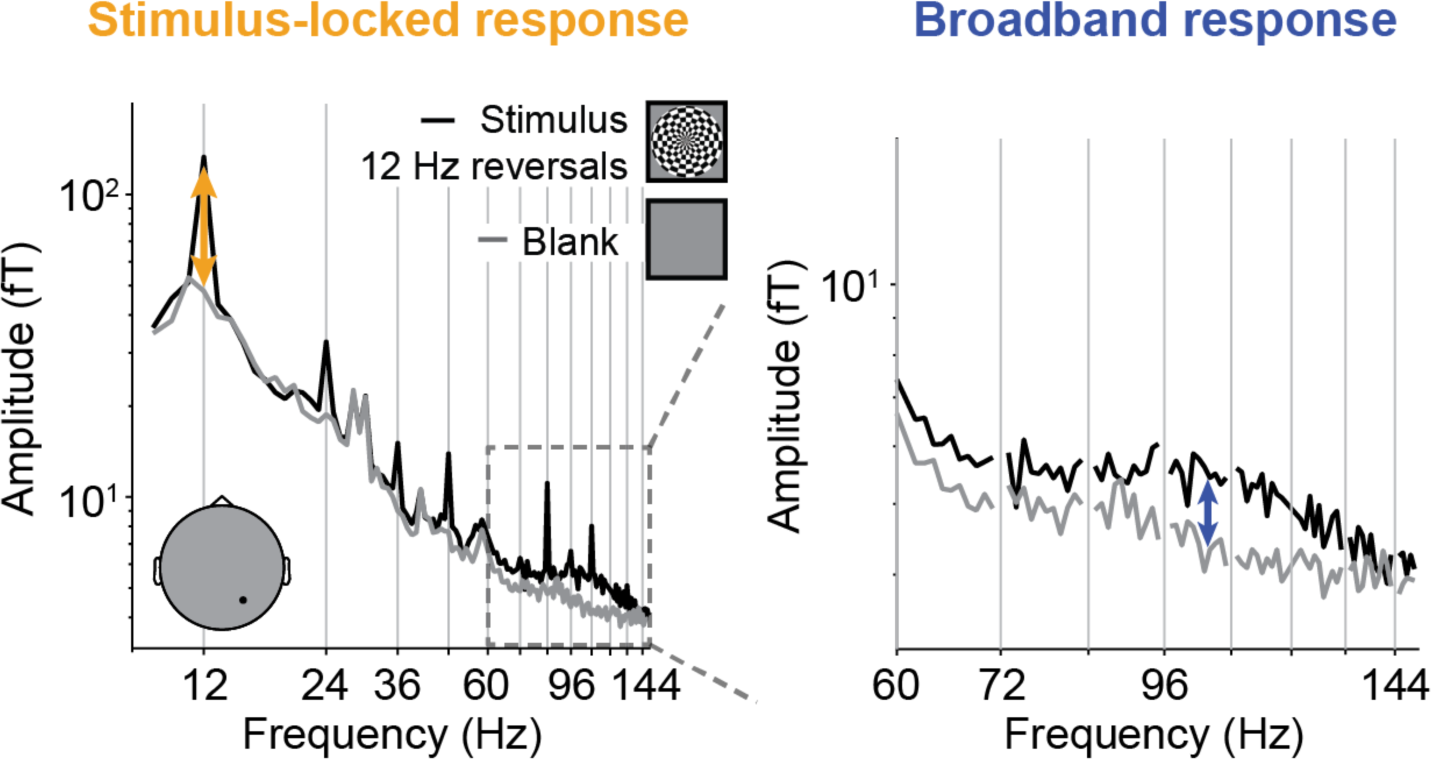
Separating the MEG response into a stimulus-locked and asynchronous component. Left panel: Average amplitude spectrum of a single posterior MEG sensor for a contrast-reversing pattern and blank (mean-luminance) periods. The black dot in the gray head schematic indicates the sensor location. The amplitude spectrum was computed in one-second epochs and averaged across ∼300 epochs per condition. The response at 12 Hz (orange arrow) and harmonics are time-locked to the stimulus, which contrast-reverses 12 times/s. **Right panel:** Zoom of frequencies from 60-150 Hz reveals a broadband elevation (blue arrow). The calculation of broadband responses excludes frequencies that are multiples of the stimulus frequency (12 Hz). Data from subject S12.

### 2.2 Stimulus-locked and broadband responses differ in their sensor topography

Both stimulus-locked and broadband responses are largest in posterior sensors, as expected from neural activity in visual cortex. However, the two components show distinct spatial topographies across these posterior sensors (**Figure 3**). The stimulus-locked response is split into two groups of sensors, extending laterally to left and right. There is a decreased amplitude in the central posterior sensors (**Figure 3**, top row). This pattern holds for individual subjects and data that are sensor-wise averaged across subjects.

**Figure 3.**
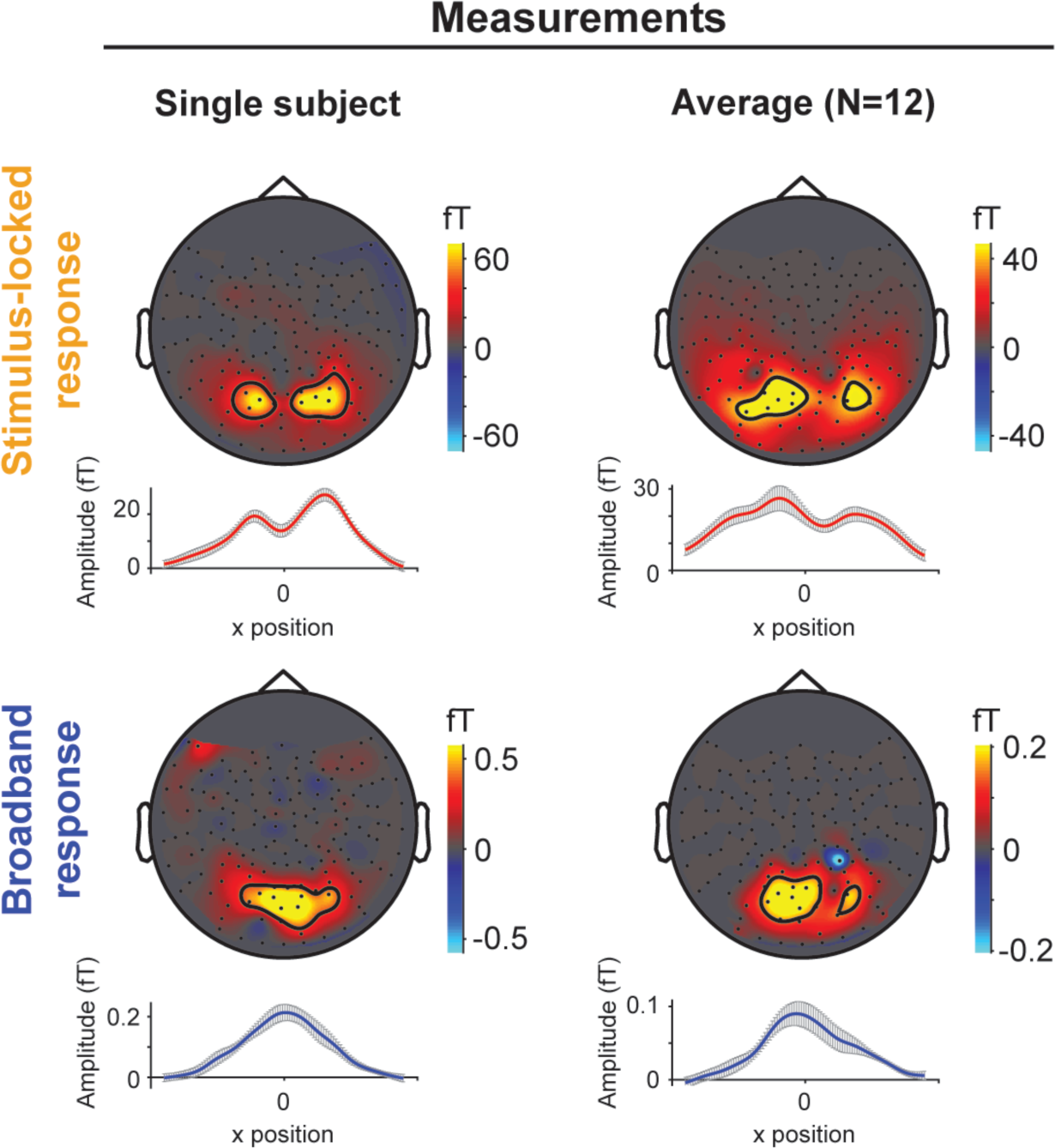
The stimulus-locked and broadband responses show different spatial topographies. **Top row:** The stimulus-locked responses on topographic MEG sensor maps and 1-dimensional summary representations of posterior sensors. The black contour lines indicate the 10 sensors with the largest response amplitudes. Amplitudes are set to 0 if the signal-to-noise ratio was below 1 (see Methods). In both single subject and group average maps, the topography shows two laterally displaced regions with large responses. Line plots show the average across posterior MEG sensors from left to right in 100 bins (red line) with gray error bars representing the standard deviation across bootstraps for the individual subject (left column) or standard error of the mean across subjects for the group average (right column). **Bottom row:** Same as top row but for broadband response. The broadband topography differs from the stimulus-locked topographies, with the largest responses in the central posterior sensors. Data from single subject S12. Group average topographic maps depicts sensor-wise averaging. (See **Supplementary Figure S1** for individual data from all subjects). For a movie of the responses unfolding over time, see *makeFigure3movie.m*.

Because we define the stimulus-locked responses as the amplitude component of the Fourier transform at the stimulus frequency, the plots do not show phase data. The phases of the left and right two regions are in approximate counter-phase in both the single subject and the group average (see the script *makeFigure3PhaseMaps.m*).

The broadband responses differ in their spatial topography: the largest response is in the central posterior sensors, at approximately the same location where the stimulus-locked responses show a decrement in amplitude (**Figure 3**, bottom row). The overall spatial pattern of broadband responses is unimodal, whereas the stimulus-locked responses tend to be bimodal. These differences in spatial topographies can be seen at both group level and in individual subjects (**Figure 3** and **Supplementary Figure S1**).

We considered whether the difference in response pattern could be due to the fact that the stimulus-locked response is defined at a single frequency, whereas the broadband response is spread across a large frequency band. For example, if the broadband response was also bimodal, but the specific pattern differed across temporal frequency bands, then combining across bands might blur out the distinct spatial peaks. We checked for this possibility by analyzing the spatial pattern of the broadband response in separate, narrow frequency bands, and find that each band tends to show unimodal patterns, similar to what we find for the analysis combined across 60-150 Hz (**Supplementary Figure S2**).

Moreover, when the stimulus-locked response is analyzed to include harmonics of 12 Hz up to 144 Hz, combining across many frequencies, it retains a bimodal distribution (**Supplementary Figure S3**). Thus, the difference in pattern does not appear to arise from the bandwidth of the signals.

Why are the spatial topographies of stimulus-locked and broadband responses different? Typically, a difference in sensor topography would be interpreted as a difference in source topography. For example, the broadband responses might arise from one set of visual areas and the stimulus-locked response from a different set. However this interpretation is unlikely given prior measurements from intracranial ECoG electrodes: both stimulus-locked and broadband responses are reliably measured from the same electrodes spanning multiple early visual areas (Winawer et al., 2013).

Instead, we hypothesize that the stimulus-locked and broadband responses measured in the MEG sensors both originate from sources in early visual cortex, differing in their temporal properties rather than spatial properties. Specifically, we speculate that the same (or similar) cortical locations generate both types of responses, but that the sources generating the stimulus-locked response are synchronized across a large spatial extent, whereas those generating the broadband response are not.

### 2.3 An MEG encoding model: predicting sensor responses from the stimulus

To assess whether a difference in temporal properties alone could account for the observed differences in spatial topographies, we simulate two types of cortical activity, one with widespread neural synchrony (*synchronous*) and one without (*asynchronous*), with both types of activity arising from the identical set of cortical locations. For the synchronous and asynchronous simulations, we separately compute the predicted spatial topographies in the MEG sensors. We then compare these predicted topographies to the observed MEG topographies from the stimulus-locked and broadband data components.

Importantly, whether or not there is widespread neural synchrony is an independent question from whether or not the responses are time-locked to the stimulus. For example, it is possible that neuronal responses contain frequencies not in the stimulus, but that are synchronized across space. These responses would be synchronous but not time-locked to the stimulus. This might occur if multiple cortical locations respond with the same temporal non-linearities. The converse is also possible: each local region could be time-locked to the stimulus, but differ in phase, rendering the responses asynchronous (out of phase) across space.

We developed an encoding model that takes a visual stimulus as input, generates a predicted response on the cortical surface, and projects these predicted cortical responses to MEG sensors. To do so, we first extract a spatial and temporal feature from the stimulus: the contrast aperture and the contrast-reversals (**Figure 4**, Step 1.1).

**Figure 4.**
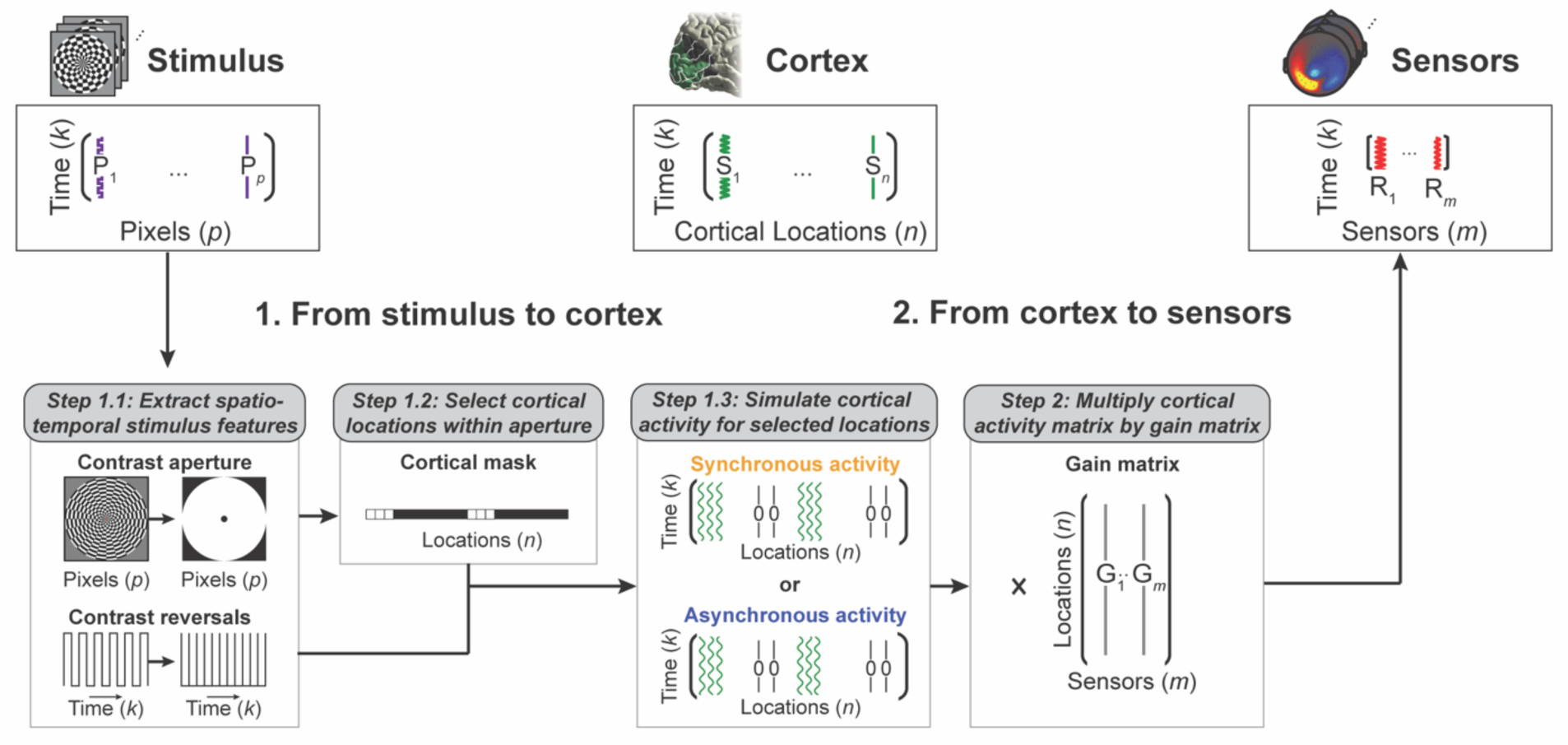
Stimulus-referred modeling approach. *From stimulus to cortex.* **Step 1.1** The high contrast-reversing dartboard pattern is reduced to two features: a spatial and a temporal feature. The spatial feature is the contrast aperture, by reducing the stimulus to a single image which will be binarized. The temporal feature is the contrast-reversal rate (12 reversals/s), by treating each pixel’s luminance change as a single contrast-reversal (thus luminance changes from black-to-white as well as white-to-black). **Step 1.2:** The contrast image is used to build a cortical mask: select cortical locations in V1-V3 which preferred population receptive field (pRF) center falls within the stimulus aperture, by applying the anatomical retinotopy template developed by Benson *et al*. (2014) to every individual’s cortical surface. **Step 1.3:** V1-V3 cortical activity is simulated as sine waves or zeros and can be described as a matrix with *k* time points by *n* locations (represented as green vertices). The degree of neural synchrony is set by a binary synchrony parameter. This parameter defines whether simulated cortical activity will be synchronous (phase-locked) or asynchronous (phase randomized) across cortical locations. In both simulations, sine waves have unit amplitude, a frequency equal to the contrast-reversal rate and identical cortical locations (*i.e.*, the only difference is their relative phase). ***From cortex to sensors.* Step 2:** To project the neural activity from cortex to MEG sensors, we multiply the cortical activity matrix by the gain matrix. This gain matrix contains a set of weights (*n* locations by *m* sensors) which defines the contribution of each source to each sensor, derived from the cortical geometry and the sensor locations. This multiplication results in predicted MEG responses (*k* time points by *m* sensors).

The contrast aperture is then used to select cortical locations (surface vertices) in visual areas V1, V2 and V3 with preferred visual field centers falling within the aperture (**Figure 4**, Step 1.2). These visual field preferences are computed for each individual subject by applying the anatomical retinotopy templates by Benson *et al*. (2014) to the reconstructed cortical surface of a T1-weighted anatomical image.

The contrast-reversals are used to simulate the neural time series, which we assume to be a harmonic at the contrast-reversal rate (**Figure 4**, Step 1.3). For all the selected surface vertices, we assume unit amplitude and a fixed frequency. Other sources (locations outside V1-V3 or with visual field locations outside the stimulus aperture) have a time series with no modulation (all zeros). For the synchronous simulation, the harmonics for all cortical locations have the same phase. For the asynchronous simulation, the phases are randomized across cortical locations. This distinction is specified by a binary synchrony parameter.

The simulated cortical activity matrix is then multiplied by the gain matrix to generate predicted sensor responses (**Figure 4**, Step 2). The gain matrix, also referred to as the ‘volume conductor model’ or ‘forward model’, describes the weighted sum of cortical locations that contribute to each MEG sensor based on the cortical geometry, the sensor locations, and the physics of magnetic fields. Because the geometry of cortex and position in the MEG differs across subjects, the gain matrix and predicted sensor maps also differ between subjects. For each subject, and on the average across subjects, we compare the predicted MEG sensor responses with the observed MEG sensor responses by summarizing the predictions as the average amplitude at the input frequency across simulated epochs.

### 2.4 Source synchrony affects predicted MEG sensor topography

Our model predictions show qualitatively different spatial topographies depending on the underlying synchrony. For synchronous sources, the model predicts the largest amplitudes in two lateralized groups of posterior sensors, separated by a decrease in amplitude in the central posterior sensors (**Figure 5**, top row). In contrast, model predictions for asynchronous sources can be characterized as a single region of large response, located at the central posterior sensors (**Figure 5**, bottom row). The differences in predicted topographic sensor maps for synchronous versus asynchronous simulations are clear at both the group level and in individual subjects. The general patterns hold for different methods used to derive the gain matrix (for predictions using the Boundary Element Model method, see **Supplementary Figure S7**).

**Figure 5.**
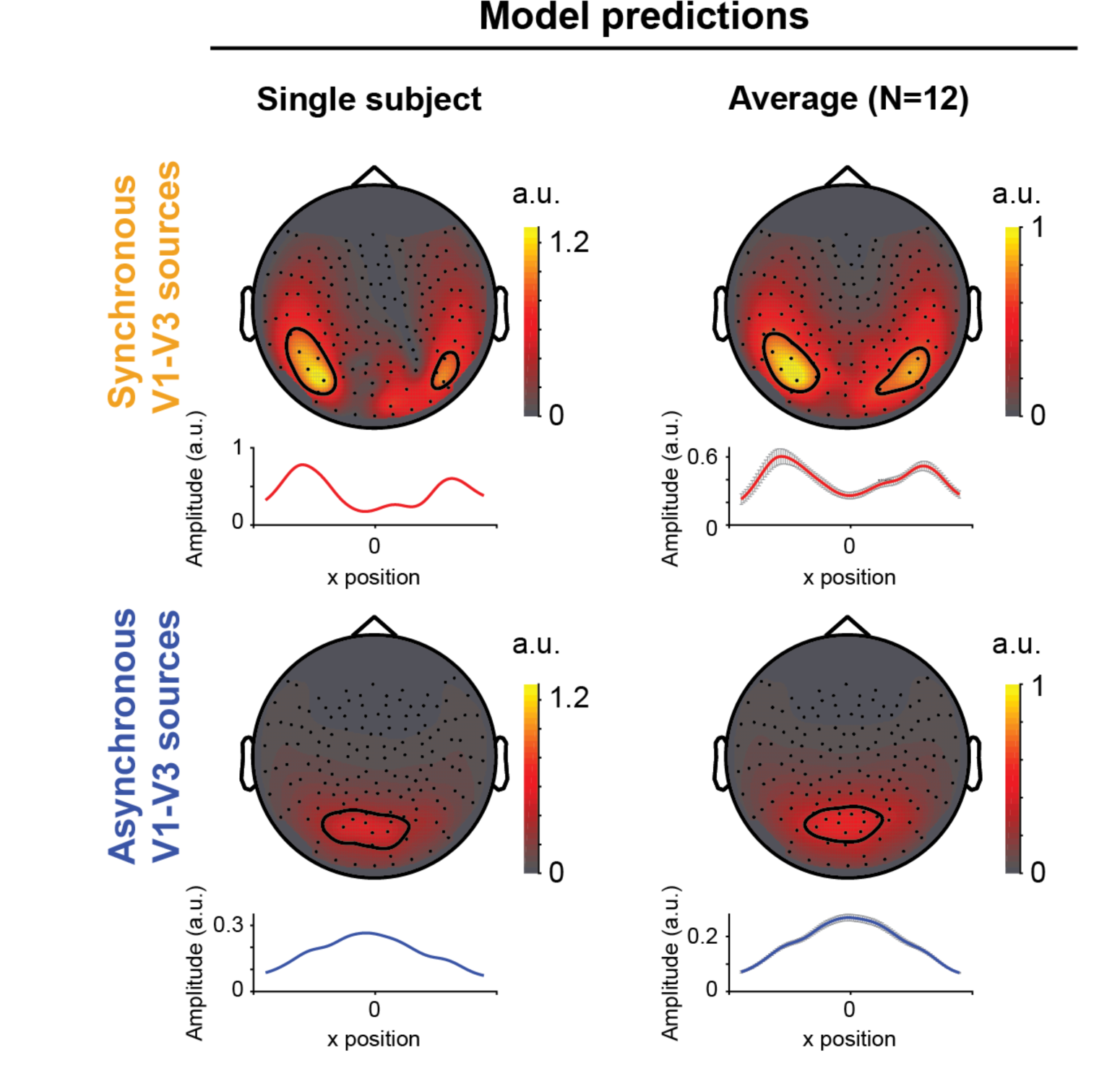
Visual encoding model predicts different spatial topography depending on synchrony, matching observed MEG data. **Left column:** Sensor-wise average for a single subject (S12). The first column shows the model output for sources that are synchronous (top) or asynchronous (bottom) across space. The synchronous sources in the model result in two laterally displaced response peaks in the sensor map and 1-dimensional summary representations of posterior sensors, as do the stimulus-locked responses in the data. The asynchronous sources in the model result in a single central response peak, approximately matching broadband responses. Predicted sensor responses for both synchronous and asynchronous simulations are normalized to the largest response in the synchronous spatial map. Line plots show weighted average across posterior MEG sensors in 100 bins from left to right. In contrast to Figure 3, individual subject’s line plots do not have error bars as the simulation does not include noise. **Right column:** Same as panel A but sensor-wise group average across all 12 subjects. For all individual subject model predictions, see **Supplementary Figure S4**. For a comparison of model predictions using only V1 or V2-V3 without V1 sources see **Supplementary Figure S5**. For a movie of the predicted responses unfolding over time, see *makeFigure5Movie.m*.

### 2.5 There are shared topographic features between the two model predictions and the two data components

The difference in spatial topography across MEG sensors between the synchronous and asynchronous model predictions bears some similarity to the observed difference between the stimulus-locked and broadband responses. First, the asynchronous simulation predicts one centralized response region, similar to the topography of broadband responses (compare **Figures 3** and **5**, bottom rows). Second, the synchronous simulation predicts two lateralized response regions, also observed in the stimulus-locked sensor topography (compare **Figures 3** and **5**, top rows). The precise locations of the lateralized regions differ between the model predictions and the stimulus-locked data, with the model outputs being more lateralized than the stimulus-locked data. There are several possibilities for this discrepancy, which we return to in the Discussion. Nonetheless, the fact that the two distinct data topographies are qualitatively predicted from the two simulations is surprising, given that the two simulations use exactly the same cortical sources, with exactly the same frequency and amplitude, differing only in their relative phase across cortical locations (single phase versus randomized phases).

The two simulations differ in predicted amplitude as well as spatial topography, with larger sensor responses for the synchronous than the asynchronous simulation. In particular, the peak response from the synchronous simulations is about 2x larger than the peak response from the asynchronous simulation. These peaks are at different sensors, with the synchronous peak lateral and the asynchronous peak more central. Comparing the same sensors within the lateral regions, the responses are about 10x larger for the synchronous than the asynchronous sources.

Differences in response amplitude are also observed for the two components of the MEG data, with the stimulus-locked responses much larger than the broadband responses. For example, for the sensor in **Figure 2**, there is an approximate 2-fold (189%) increase in stimulus-locked amplitude and only a 14% increase in broadband amplitudes, consistent with our previous observations comparing these two signal components (Kupers et al., 2018).

This observed amplitude difference is in line with the hypothesis that stimulus-locked responses arise from widespread synchronous cortical activity while the broadband responses arise from largely phase-randomized activity: The sum of synchronous activity will tend to be larger than the sum of asynchronous activity (Krusienski, McFarland, & Wolpaw, 2012; Winawer et al., 2013; Hermes et al., 2017; Kupers et al., 2018).

### 2.6 Cancellation of synchronous source activity explains lateral sensor topography predicted by encoding model

Why do the model predictions from synchronous neural sources show bimodal activation peaks in the sensor topography, whereas predictions from asynchronous sources do not? One possibility is that the predicted ‘gap’ between the two activation peaks for synchronous sources (but not asynchronous sources) is caused by large-scale cancellation of neuroelectric fields from opposite-facing dipoles, arising from the cortical geometry (folding pattern). Occipital cortex is highly folded (Duvernoy, 1999) and each of the early visual maps, V1-V3, contain deep sulci. As a result, a large stimulus such as the one in our experiments, will activate a broad swath of these maps, resulting in many locations where there are opposite-facing dipoles from either side of the sulcus. For example, in Subject S12, there are large regions in each of V1, V2, and V3 which include two opposing sulcal banks (**Figure 6**). Because multiple visual field maps could contribute to signal cancellation, we consider the effects of simulations with only subsets of the maps (**Supplementary Figure S5**). These show that simulations with V2 and V3 (without V1) tend to show model predictions similar to those that include V1-V3, indicating a likely contribution from V2 and V3 in terms of large-scale signal cancellation, consistent with previous analyses (Ales, Yates, & Norcia, 2010).

**Figure 6.**
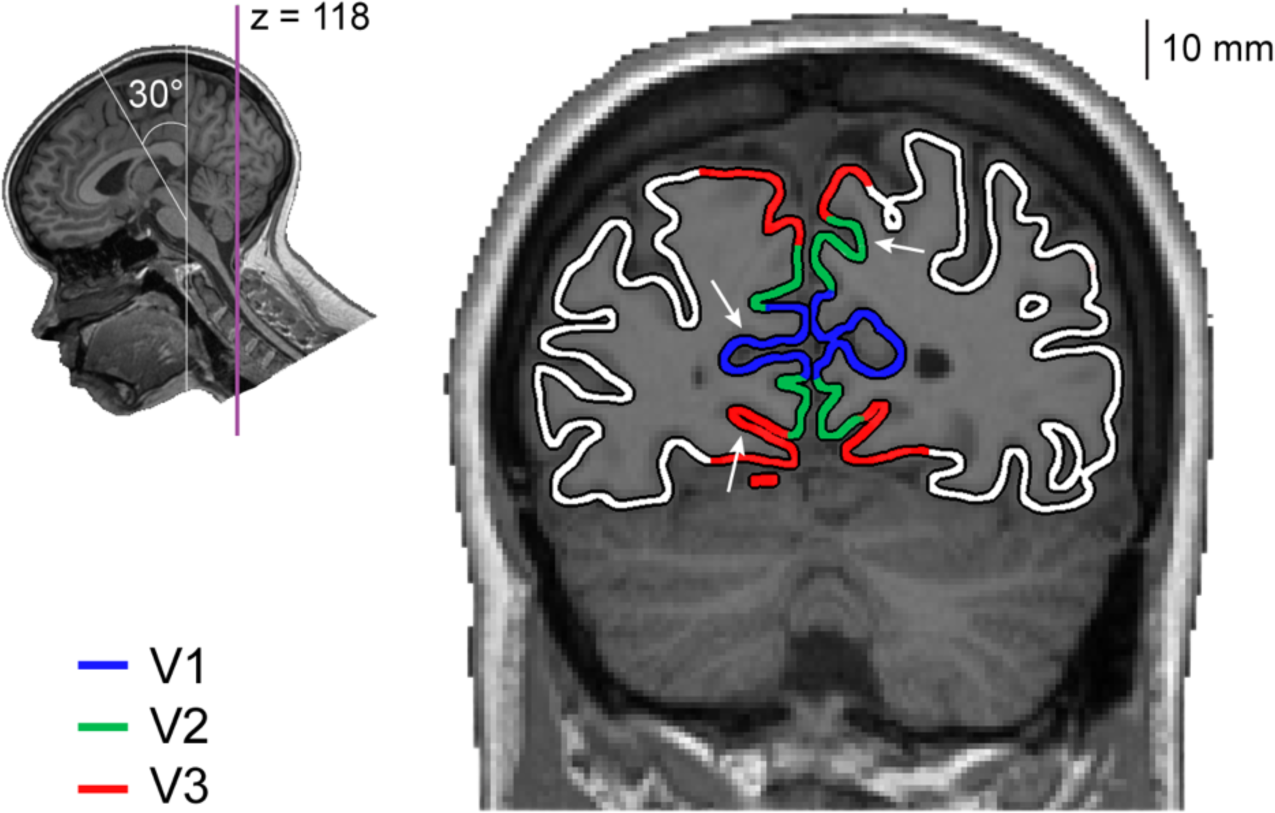
An example folding pattern in occipital cortex. The figure shows an oblique slice perpendicular to the calcarine sulcus (purple line) from a T1-weigthed image of subject S12. In each of the V1-V3 maps, there are regions containing opposite-facing sulcal walls. Three examples are indicated by the white arrows.

The cortical folding pattern interacts with synchronous and asynchronous sources differently. Synchronous source activity will add when the dipoles are parallel, causing a large response at the sensor level, and cancel when the dipoles are opposite facing (**Figure 7**, top row). For asynchronous sources, there is partial cancellation regardless of whether the dipoles are parallel or opposite facing (**Figure 7**, bottom row). Overall, the largest sensor signals will come from synchronous sources that are parallel, with lower signal from asynchronous sources (**Figure 7**, upper left vs lower left).

**Figure 7.**
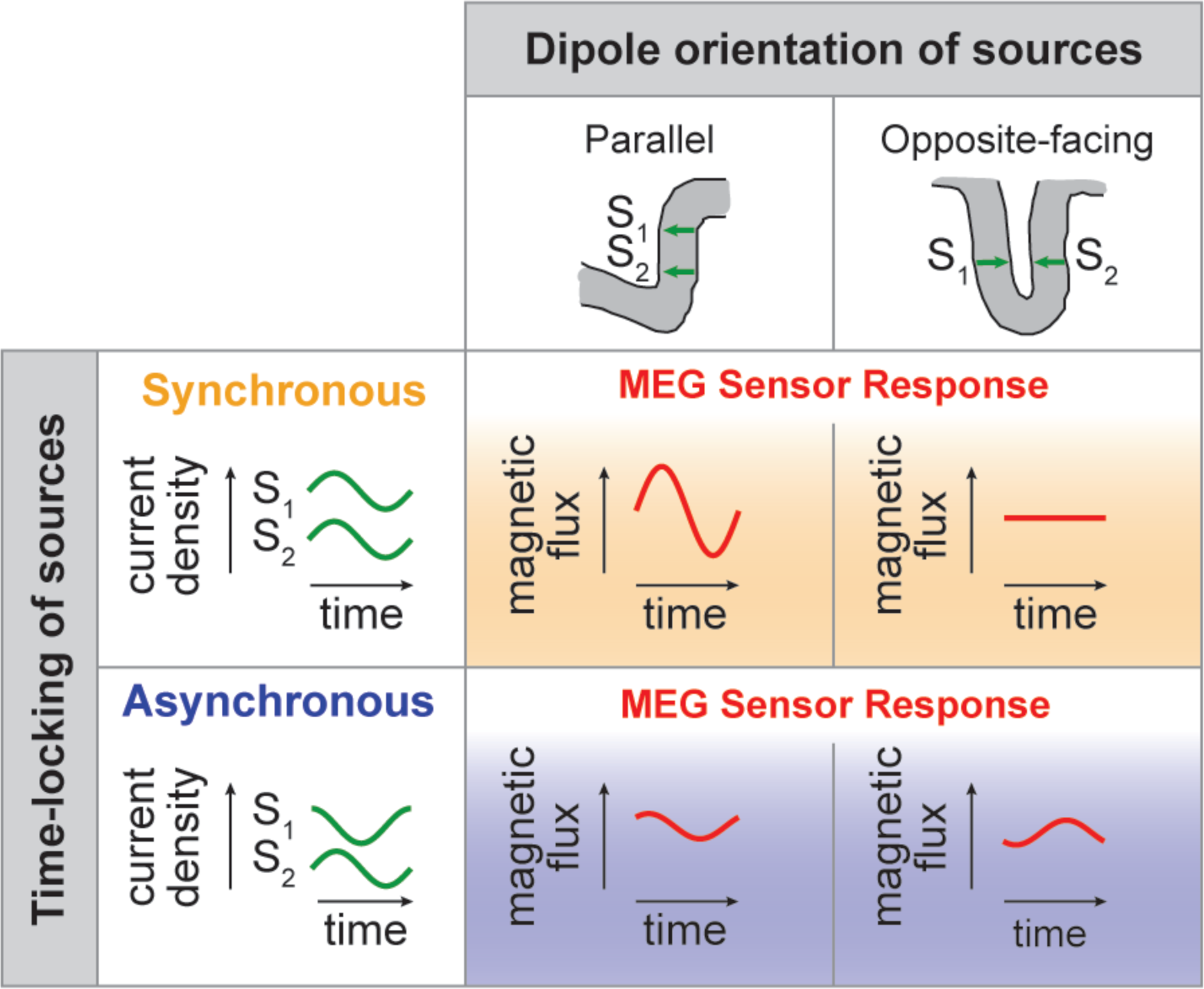
The effect of cortical geometry and neural synchrony on MEG sensor responses. The cancellation hypothesis predicts that a difference in spatial topographies across MEG sensors is caused by an interaction between the geometry of the cortex and the degree of neural synchrony. **First column:** If neighboring sources (S1 and S2, green sine waves) are located such that their dipole orientations are parallel to each other, their responses will add at the sensor level. **Second column:** In contrast, if sources are located such that their dipole orientations are opposite-facing their source responses will interfere at the sensor level. Besides the relative dipole orientation, the summation or cancellation of underlying source response also depends on the time-locking (or relative phase) of sources contributing to the MEG sensor responses (red sine waves): synchronous sources will largely sum or cancel at the sensor level (first row, yellow gradient), whereas asynchronous sources will partially cancel at the sensor level (second row, blue gradient).

To test whether the bimodal distribution in sensor topography from the synchronous simulation is caused by cancellation arising from the cortical geometry, we modified our encoding model by making the gain matrix positive only. A positive-only gain matrix preserves the sign of the source activity when projected to the sensors so that synchronized source activity cannot cancel. Changing the gain matrix in this way has a big effect on the sensor predictions from synchronous sources (**Figure 8**, top row), but not asynchronous sources (**Figure 8**, bottom row). For this positive-only gain matrix, the difference in topography between the synchronous and asynchronous simulations disappears (**Figure 8**, 2^nd^ column). This topography closely resembles the observed topographic maps for the broadband component of the MEG responses (**Figure 3**, bottom row). These results indicate that the idiosyncratic ‘gap’ in the spatial pattern of the synchronous simulation is caused by cancellation of neuroelectric fields, presumably from opposite facing dipoles in upper and lower bank of the Calcarine sulcus. When preserving the sign of dipoles our encoding model, we showed that spatial topography predicted by synchronous sources is caused by signal cancellation due to the cortical folding in early visual cortex.

**Figure 8.**
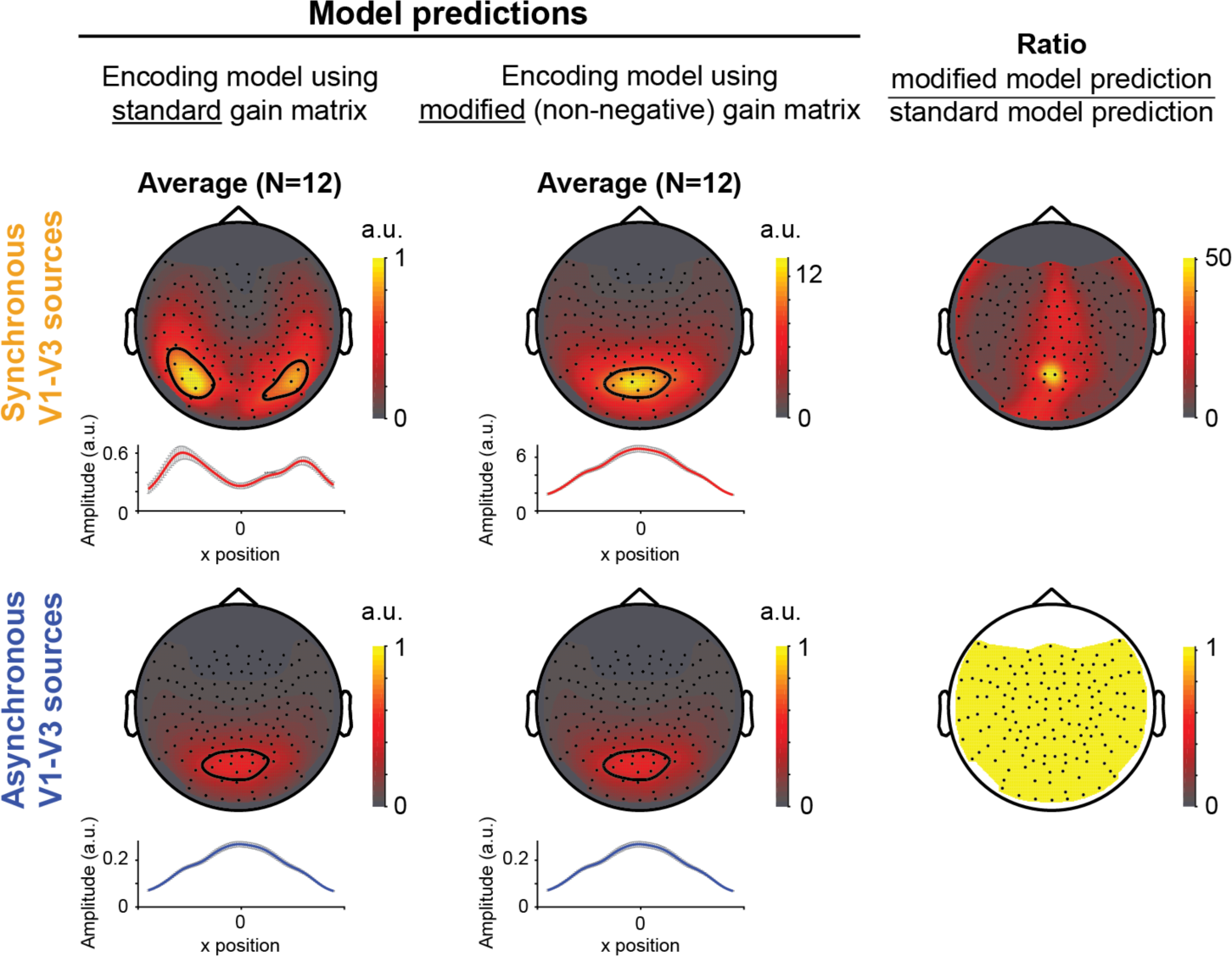
Preventing cancellation in visual encoding model affects predicted MEG sensor responses for synchronous, but not asynchronous sources in early visual cortex. **First column:** Predicted MEG sensor amplitudes using a standard forward model which allows for neural sources to cancel their current density. Data are identical to first column of Figure 5. **Second column:** Predicted sensor amplitudes using modified forward model, *i.e.*, using the absolute values of gain matrix, preventing neural sources to cancel their current density. Data are plotted to with the same scaling as in the left column. **Third column:** Ratio of predicted sensor amplitudes predict by the standard and modified encoding model (modified model divided by standard model). Preventing cancellation affects the topography of synchronous, but not asynchronous of V1-V3 sources. The contour lines are drawn at 93.6^th^ percentile of the model predictions, corresponding to the 10 sensors with the largest response. Data are from group average.

## 3. Discussion

We developed a visual encoding model for MEG that predicts sensor responses to stimuli and showed that the predicted topographies in MEG sensors depend on whether cortical responses are synchronous or asynchronous. We compared these model predictions to observed MEG data from subjects viewing a contrast-reversing pattern separated into a stimulus-locked and a broadband component. We found that the two data components have different spatial topographies in the MEG sensors, where the topography of the stimulus-locked data component was similar to the synchronous simulation, and the topography of the broadband data component was similar to the asynchronous simulation.

### 3.1 Cortical geometry mediates the relationship between source synchrony and sensor topography

The simulations support the interpretation that the differences in spatial topography between the two data components lie in the temporal properties of source activity, not spatial properties. To understand how temporal properties of the neural responses influence spatial topography in the sensor data, it is necessary to consider the cortical geometry.

Previous studies have argued that the geometry of primary visual cortex results in a peculiar property of the V1-driven evoked (time-locked) EEG response. Specifically, according to ‘the cruciform hypothesis’, responses to stimuli in the lower visual field and upper field can result in a similar voltage time series except with opposite polarity (Jeffreys, 1971; Jeffreys & Axford, 1972b, 1972a). This result has been attributed to the fact that the upper and lower field representations of V1 lie on opposite sides of the Calcarine sulcus, resulting in opposing dipoles. This explanation is consistent with our observation that the time-locked portion of the MEG response to a large stimulus (containing contrast in both lower and upper visual field) tends to result in a spatial gap in the topography: the gap is where signals from opposite-facing dipoles cancel. This interpretation is supported by our simulations showing that when the effect of dipole cancellation is removed, there is no gap (**Figure 7**).

Related questions have received considerable interest in recent years: namely, whether the upper and lower field representations of V1 perfectly cancel in EEG sensor responses, and if V1 is the only visual area that can exhibit this phenomenon (Ales, Yates, et al., 2010; Ales, Yates, & Norcia, 2013; Kelly, Schroeder, & Lalor, 2013; Kelly, Vanegas, Schroeder, & Lalor, 2013). Our observation that the stimulus-locked signal has a systematically different spatial topography than the broadband signal does not depend on whether the cancellation comes from V1 only, or also from V2 and V3. In fact, sensor cancellation in both EEG and MEG can result from spatially extended source activity in any cortical location, or even from small patches of randomly distributed responses across cortex (Ahlfors et al., 2010). The question of whether synchronous source activity causes cancellation at the sensor level depends only on whether opposing dipoles are simultaneously activated. Since there are more likely to be opposing dipoles in areas of cortex with high curvature, there is a relationship between the cortical curvature, neural synchrony, and the spatial pattern in the sensor responses.

The prior studies focused on evoked potentials (Ales, Yates, et al., 2010; Ales et al., 2013; Kelly, Schroeder, et al., 2013; Kelly, Vanegas, et al., 2013) and evoked fields (Ahlfors et al., 2010), defined by trial-averaging the data time-locked to stimulus onset, or simulating simultaneous sources. A novel finding in this study is that the broadband signal does *not* show cancellation and is thus much less affected by the details of the cortical geometry. This is evident in both data and simulations. We conclude from this that the broadband signal is asynchronous not only with respect to the stimulus, which is true by definition of this data component, but also across space.

Our modeling was motivated by the observation that the spatial topography differed between the stimulus-locked and broadband responses. The particular feature that stood out was the bimodal spatial pattern in the stimulus-locked response. This pattern appears in a number of other studies employing visual steady paradigms (Moratti, Keil, & Miller, 2006; Kamphuisen, Bauer, & van Ee, 2008; Giani et al., 2012; Pisarchik, Chholak, & Hramov, 2019) or in the evoked response to presentation of static images (Mecklinger et al., 1998; Schoenfeld, Heinze, & Woldorff, 2002; Golubic et al., 2011). Not every study observes this bimodal pattern (*e.g.* see figure 2C in (Zhigalov, Herring, Herpers, Bergmann, & Jensen, 2019), which does not show this pattern). Our proposal that the bimodal pattern in our study reflects signal cancellation may also explain the pattern in some of these other studies. However, the predicted spatial pattern in the sensors is highly dependent on features of the individual subject’s cortical and head geometry, as well as on which sources are most active. For example, we see that simulations that omit V1 sources still predict bimodal distributions, whereas simulations that omit V2/V3 do not (**Supplementary Figure S5**). Hence, for any particular study, whether or not the sensor pattern looks bimodal will likely depend on spatiotemporal properties of the cortical response, which are stimulus dependent, the type of sensors, and the geometry of the individual’s head and brain.

### 3.2 Asynchrony between cortical sources reduces the amplitude of sensor responses

As discussed in the previous section, synchronous cortical activity results in cancellation due to the cortical geometry. This influences the sensor spatial topography. Asynchronous cortical activity also results in cancellation, but for a different reason and at a different spatial scale. The asynchrony in the neural responses means that their time series may be rising in one cortical location while falling in a neighboring location. This asynchrony may play out at a very fine scale, *e.g.* < 1 mm, perhaps within single cortical columns. This kind of local cancellation does not depend on the cortical folding pattern, which varies slowly at the sub-mm scale. Hence, asynchrony between sources has a general effect of reducing the amplitude across the sensor array. At the extreme, the summed response of *n* cortical sources with random phases will grow with the square root of *n*, and with synchronous sources with *n.* As a result, we expect synchronous source activity to translate to large sensor responses. This is confirmed in our simulations in which equal amplitude neural responses result in large or small sensor responses. This logic has been used to explain why both evoked and oscillatory ECoG responses are large compared to broadband responses (Winawer et al., 2013; Hermes et al., 2015; Hermes et al., 2017). The same principle can also explain why scalp EEG responses at lower temporal frequencies are large: lower temporal frequencies are more synchronous across space than higher frequencies (Pfurtscheller & Cooper, 1975).

In sum, our model captures two types of cancellation. One type is the cancellation from synchronous activity across large-scale variation in cortical geometry (such as the folding of the large sulci in visual cortex), which translates to a spatial effect at the sensor level. The second type of cancellation arises from asynchronous activity in local responses and causes an overall amplitude reduction across the sensor array.

This second type of cancellation has an important implication for the interpretation of neuroscience data: the largest signal measured by the instrument cannot be assumed to reflect the largest amount of underlying neural activity (see similar reasoning in (Musall et al., 2014; Butler, Bernier, Lefebvre, Gilbert, & Whittingstall, 2017; Hermes et al., 2017). For the same reason, evoked potentials, although they can be quite large, are often a poor predictor of the fMRI BOLD signal, which is relatively insensitive to neural synchrony at the millisecond scale (Foucher, Otzenberger, & Gounot, 2003; Winawer et al., 2013).

### 3.3 Phase-locking across cortex vs time-locking to the stimulus

One of the interesting observations from our simulations is the qualitative match between the synchronous simulation and stimulus-locked data on the one hand, and the asynchronous simulation and broadband data on the other hand. Our results here suggest that the time-locked response to our contrast-reversing stimuli is in large part also synchronous across space, whereas the broadband response is not synchronous across space. This pattern may be a general feature of visual encoding in early visual areas. A rapid set of coherent signals arrive in visual cortex, giving rise to the time-locked signal. These responses then set off a cascade of local intracortical process, which are not synchronized across space or to the stimulus. We described these as two visual circuits in prior work (Winawer et al., 2013).

One might be tempted to reason that our results are circular. If the signals are time-locked to the stimulus, are they not necessarily synchronized? And if they are asynchronous with one another, is it not the case that they will necessarily be asynchronous with respect to the stimulus? In fact, this is not correct. The agreement between data and simulations are not a consequence of definitions, but rather they are an empirical result. For example, it is possible for cortical responses to be synchronized *across space* but *not time-locked* to a stimulus onset. This occurs, for example, with certain pharmacological manipulations that increase long-range synchrony even without a stimulus (Musall et al., 2014). Increased synchrony across space that is not time-locked to stimulus onset also occurs with gamma-band oscillations in response to certain types of stimuli. These spatially synchronous but non-time locked responses are sometimes called ‘induced’ rather than ‘evoked’ oscillations (Tallon-Baudry & Bertrand, 1999). Oscillations that are thought to be synchronous across space but not time locked to a stimulus also occur in other frequency bands, such as alpha and beta bands (Berger, 1929; Adrian & Matthews, 1934).

The converse is also true: neural responses can be stimulus-locked but not synchronous across space. For example, recordings across cortical depth show that there are stimulus-locked (‘evoked’) responses whose phase varies across depth, sometimes resulting in a phase reversal between supragranular and infragranular layers (Haegens et al., 2015). Even though these responses are out of phase with each other, they will each show their own time-locked response to stimulus onset. Similarly, responses in different cortical regions (say V1 and parietal cortex) may each be time-locked to the stimulus but vary in onset (hence asynchronous across space) (Chen et al., 2007; Cottereau et al., 2011). Hence whether a response is time-locked to a stimulus or not is independent from whether it is highly synchronous across space.

### 3.4 Where does the broadband response come from?

In our forward model, we used a traditional approach of minimizing differences between conditions: The synchronous and asynchronous simulations were identical in all ways except one (the degree of phase-locking between cortical locations). An alternative approach would be to use a more biophysical simulation, including model neurons that generate both evoked and asynchronous responses. To do so requires a biophysical model that generates high frequency broadband responses. There are several models of how this might arise, including the temporal integration of Poisson spike arrivals in synapses and dendrites (Miller et al., 2009), and the tendency for neurons to exhibit up-down phase changes (Milstein et al., 2009). Another possibility is that responses at non-stimulus frequencies arise from non-linear interactions between neural responses at the driven frequency and the background neural activity. Such nonlinearities have been postulated to explain variability in the EEG responses at the stimulus frequency, and could also contribute to responses outside the driven frequency (Mast & Victor, 1991; Victor & Mast, 1991). Following our prior work (Winawer et al., 2013), we implemented model neurons adapted from the Miller model, and combined their outputs with the MEG head model (gain matrix) to predict time series in the MEG sensors. This more biophysically realistic simulation produces the same pattern of effects as the simpler simulation: a unimodal spatial distribution of sensor responses for the broadband signal, and a more bimodal distribution for the stimulus-locked (**Supplementary Figure S6**).

Alternatively, in principle, it is possible that broadband responses measured extracranially (EEG or MEG) could arise from non-neural sources, such as artifacts from eye movements (Yuval-Greenberg, Tomer, Keren, Nelken, & Deouell, 2008), head muscle contractions (Muthukumaraswamy, 2013), or environmental electromagnetic noise. We don’t believe these artifacts explain the broadband responses we observed. First, as we demonstrated in our previous paper measuring broadband responses (Kupers et al., 2018), neither the rate nor the direction of microsaccades varies systematically with the observed broadband responses in individual subjects: Subject data containing the highest microsaccade rate do not show the largest broadband response and vice versa, and the direction of microsaccades does not systematically bias to left or right visual field.

Moreover, when removing epochs with microsaccades from the analysis, the broadband response remained evident in each individual subject.

Second, the characteristics of these noise artifacts do not overlap with our measured broadband responses (see also the Discussion section “Challenges in measuring extracranial broadband responses” in (Kupers et al., 2018)). For instance, the MEG spike field artifact, which can be confused with broadband neural activity (Yuval-Greenberg & Deouell, 2009), has a spatial topography affecting mostly temporal and frontal MEG sensors (Carl, Acik, Konig, Engel, & Hipp, 2012). Our broadband responses are confined to middle posterior sensors and therefore are unlikely to be contaminated by spike field artifacts.

Saccadic spike potentials, caused by the retina-to-cornea dipole, introduce artifacts between 4-20 Hz (Keren, Yuval-Greenberg, & Deouell, 2010). These frequencies are lower than the range we use to define our measure of broadband responses (60-150 Hz), hence unlikely to affect our measurements. Although, head muscle contraction and environmental noise contributions have been reported to cause spectrally broad artifacts (Muthukumaraswamy, 2013), it is unlikely that these noise sources only affect central occipital sensors but not lateral occipital MEG sensors.

### 3.5 Relationship to previous work on modeling MEG/EEG signals

MEG/EEG data have been subject to a variety of computational models, which can broadly be divided in three groups. The approaches with the longest history and most used are forward models (predicting sensor responses from cortical data) and inverse modeling (predicting cortical responses from sensor data). In addition, decoding models have started to become more widely used. The models infer the stimulus (or stimulus features) from the sensor data. From the three groups, our visual encoding model overlaps mostly with the forward modeling approach, but differs in that it predicts the sensor responses from stimulus features.

#### 3.5.1 Inverse and constrained inverse models

Inverse models are used to infer the cortical source activity from the MEG/EEG sensor responses. They neither predict brain activity nor sensor responses from the input (*e.g.*, a visual stimulus), differing from our approach here. A benefit of these models is that no *a priori* knowledge of neural sources is needed. However, inverse models are ill-posed, since there are many more cortical sources than there are MEG or EEG measurement channels (Hämäläinen et al., 1993). To the degree that source activity differs from model assumptions, it will be inaccurate. Consider our two simulations (synchronous and asynchronous); they had activity in the identical set of sources, yet resulted in different spatial topographies. The correct source localization would return the identical sources.

But because the inverse problem is ill-posed, this solution is not likely to be found. Instead, inverse solutions to the simulated sensor responses would incorrectly return different source locations.

Several studies have used inverse models to reconstruct sources of retinotopic responses (Moradi et al., 2003; Poghosyan & Ioannides, 2007; Sharon, Hamalainen, Tootell, Halgren, & Belliveau, 2007; Brookes et al., 2010; Cicmil, Bridge, Parker, Woolrich, & Krug, 2014; Nasiotis, Clavagnier, Baillet, & Pack, 2017). These reconstructions were able to capture retinotopic maps in early visual at a coarse scale (order of centimeters), but contained large errors (*e.g.*, failure to localize sources from the upper visual field due to penalizing the reconstruction of two active sources canceling out at the sensor level).

One approach to improve localization error is Retinotopy Constrained Source Estimation. This method uses visual field maps to guide constraints on the number of solutions when inverse modeling the sources. For example, one can create a correlation matrix that only includes visual field areas to constrain the possible solutions (Hagler et al., 2009; Hagler & Dale, 2013; Hagler, 2014; Cottereau, Ales, & Norcia, 2015) or apply an exhaustive search to define neighboring sources for every stimulus location (Ales, Carney, & Klein, 2010; Inverso, Goh, Henriksson, Vanni, & James, 2016). This approach has been shown to increase source localization accuracy, *e.g.*, decrease cross-talk for sources in visual areas with close proximity like V1 and V2. This kind of approach is complementary to ours. It is useful to infer neural responses in paradigms where an encoding model is not feasible.

#### 3.5.2 Decoding models

MEG-based decoding models have been used to predict stimulus features from sensor data, similar to the fMRI decoding literature (Haxby et al., 2001). There are many purposes for decoding models. One is to facilitate comparison with other data types when a linking model is lacking, for example comparing MEG to fMRI (Cichy, Pantazis, & Oliva, 2016) or to a neural network (Cichy, Khosla, Pantazis, Torralba, & Oliva, 2016). A second purpose is to ask how stimulus representations unfold over time (King & Dehaene, 2014), or how they are localized in temporal frequency bands (Pantazis et al., 2018). Generally, decoding models can reveal the presence of information about a stimulus or stimulus feature. In contrast, encoding models explicitly postulate computations or representations of the system, and thus offer more general system descriptions (Naselaris, Kay, Nishimoto, & Gallant, 2011).

#### 3.5.3 Forward models

Forward models compute the propagation of source activity to sensors. The source activity can either be simulated or defined by a separately acquired fMRI session (*e.g*., a retinotopy experiment), before projected from cortex to sensors. This type of computational model is closest to our approach, but again, differs from our encoding model in that these forward models do not take visual stimuli as an input.

Previous forward models have been used to create simulated sensor responses as a benchmark to test specific analyses methods, *e.g.* brain connectivity analyses in EEG (Haufe & Ewald, 2019), accuracy of volume conduction head models (Henson, Mattout, Phillips, & Friston, 2009; Stenroos, Hunold, & Haueisen, 2014; Stenroos & Nummenmaa, 2016), guide subdural electrode placement for epilepsy monitoring (Lopes et al., 2020), or provide ‘ground truth’ sources when combined with inverse modeling in both healthy (Sharon et al., 2007; Akalin Acar & Makeig, 2013; Nasiotis et al., 2017) and patient populations (Acar et al., 2008). Some recent publications of open-source EEG/MEG toolboxes have now advanced to simulating electromagnetic fields with biologically plausible noise (Barzegaran, Bosse, & Norcia, 2019) or cellular-level circuits (Neymotin et al., 2020).

In contrast to work in single-unit electrophysiology (Simoncelli & Heeger, 1998; Rust, Schwartz, Movshon, & Simoncelli, 2005; Mante, Bonin, & Carandini, 2008), functional MRI (Dumoulin & Wandell, 2008; Kay, Naselaris, Prenger, & Gallant, 2008; Kay, Winawer, Rokem, Mezer, & Wandell, 2013) and ECoG (Hermes, Petridou, Kay, & Winawer, 2019), visual encoding models that take stimuli as input and predict the measured response as output are not commonly used in EEG and MEG. This modeling approach has been highly successful in elucidating fundamental properties of sensory encoding, such as linear filtering (Enroth-Cugell & Robson, 1966; Movshon, Thompson, & Tolhurst, 1978), spatial pooling (Kay, Winawer, Mezer, & Wandell, 2013), and normalization (Heeger, 1992). Our work is one of the first to extend this approach to MEG and was useful in characterizing two types of visual responses, one highly synchronous across space, one largely asynchronous.

### 3.6 Assumptions of the model

Our visual encoding model contained two parts: an encoding model from stimulus to cortex based on retinotopic atlases (Benson et al., 2014) and a physics-based model of the propagation of cortical currents to magnetic flux at the sensors, based on overlapping spheres (Huang, Mosher, & Leahy, 1999; Tadel, Baillet, Mosher, Pantazis, & Leahy, 2011).

Both of these model components include a number of simplifying assumptions.

Advantages of the simplifications are that the model is easy to use, and when the model makes interesting predictions, they are interpretable. For instance, there were two interesting predictions made from our simulations. First, the synchronous and asynchronous simulations led to different spatial topographies (more bimodal for the synchronous simulation, and unimodal for the asynchronous simulation). This phenomenon was explained by cancellation due to the cortical folding pattern in the synchronous simulations. Second, the amplitude was generally lower for the asynchronous than the synchronous simulation. This was explained by partial cancellation between nearby sources in the asynchronous simulations. These two features of the simulations were also found in the data. Therefore, the benefit of model interpretability transfers to the data, providing plausible explanations for the phenomena observed in the experiments.

Nonetheless, it is important to consider how robust the patterns found in the simulations are: specifically, are the patterns highly dependent on the simplifying assumptions of the model?

#### 3.6.1 Encoding model assumptions

The encoding model from stimulus to cortex has three simplifying assumptions. First, cortical activity was confined to only three visual areas, V1, V2, and V3. Second, within these visual areas, responses were simulated as having the same amplitude or zero. Third, the phases of the cortical responses were either all identical (synchronous simulation) or completely random (asynchronous simulation). None of these assumptions holds exactly in the brain. For example, there are differences in EEG sensor response timing and amplitudes for checkerboard stimuli that vary in eccentricity (Jeffreys, 1971; Ales et al., 2013; Inverso et al., 2016), and higher level visual areas beyond V1-V3 can also show steady state responses to contrast-reversing stimuli (Ales, Farzin, Rossion, & Norcia, 2012). The steady state response to contrast-reversing patterns may also differ in amplitude between visual areas, as suggested by fMRI-constrained source localization of EEG signals (Di Russo et al., 2005; Di Russo et al., 2007).

In additional simulations, we tested whether deviations from these assumptions had a great impact on the main findings and found that they did not. Most importantly, we found that the difference in spatial topography —more bimodal for synchronous simulations and unimodal for asynchronous simulations— was found across a variety of simulation conditions. First, we simulated responses in 9 visual areas beyond V1-V3, as described in Benson and Winawer (2018). These simulations showed broadly similar patterns to those from only V1-V3 (**Supplementary Figure S7**). Second, we simulated responses limited to different eccentricity bands. Except when responses were confined only to the fovea, simulations showed the same general patterns as those including cortical activity for the entire stimulus aperture (**Supplementary Figure S8**). Third, we simulated intermediate levels of synchrony between 0% and 100% (**Supplementary Figure S9**).

These simulations show that fully asynchronous (0%) and fully synchronous (100%) levels are not special cases: rather, the two types of spatial patterns we found using those extreme levels were each found over a range of synchrony values. Together, these additional tests show that the pattern of results is not highly dependent on any one of the simplifying assumptions of the encoding model.

#### 3.6.2 Physics-based forward model assumptions

We used the overlapping spheres (OS) volume conductor head model to compute the gain matrix for each individual subject (Huang et al., 1999). Some investigators have argued that a model derived from a three-shells boundary element model (BEM) (Kybic et al., 2005; Gramfort, Papadopoulo, Olivi, & Clerc, 2010), while more computationally intensive to compute, is more accurate (Henson et al., 2009; Stenroos et al., 2014). To test whether our conclusions depended on using a particular forward model, we implemented the BEM model in each of our participants and found that the predictions did not differ substantially in spatial topography from the OS model for either synchronous or asynchronous sources (**Supplementary Figure S7**, panel A and B). This provides some assurance that the pattern of results obtained from the simulations are not an artifact of idiosyncratic assumptions made in the OS model. For both the overlapping spheres and the boundary element model, we assumed that the neural generators are dipoles oriented normally to the cortical surface. Some researchers have suggested that, at least in the case of EEG signals, monopole neural generators play a larger role than is traditionally thought (Riera et al., 2012), and that therefore signal cancellation from opposite-facing dipoles might have only a small influence on EEG sensor measures (Butler et al., 2019). If correct, this might explain some of the differences found between EEG and MEG measures of visually driven signals, but would not explain MEG results (Riera et al., 2012).

### 3.7 Fitting Model Parameters

We compared patterns found in our simulations to patterns observed in the MEG data, but we did not attempt to fit parameters of the model. For example, we simulated 0% and 100% neural synchrony, but did not fit a free parameter with intermediate values to best explain the data. Nor did we try to estimate the degree to which cortical response amplitude varies with retinotopic location or visual field map. To do so would require not just comparing simulations to the data, but instead specifying and fitting free parameters in a model from stimulus to cortex to sensors.

If the model has too many parameters (for example, a unique amplitude for every surface vertex), then the model will be ill-posed, similar to the problem of source localization (Hämäläinen et al., 1993; Helmholtz, 1853). Clearly, if each of the several hundred vertices in the V1-V3 maps are independently fit with one or more parameters (such as a gain factor, receptive field location, receptive field size, etc.), the problem will be ill-posed. It might be possible, however, to find a low-dimensional parameterization of the multi-variate response, making the problem well-posed. To our knowledge, a low parameter encoding model of signals across entire visual field maps does not yet exist, but there are promising developments. For example, Benson and colleagues (Benson, Broderick, Müller, & Winawer, 2017) reparametrized spatial encoding models for fMRI developed by Kay *et al*. (Kay et al., 2008; Kay, Winawer, Rokem, et al., 2013) with just a handful of global parameters.

While we reported qualitative similarities between simulations and data, we also observed quantitative differences. For example, the synchronous simulation predicts two lateralized responses which are more laterally spaced than the stimulus-locked responses. Such quantitative differences are not surprising because we did not fit model parameters to the data, as discussed above. It is likely that visual field maps beyond V1-V3 contribute to the measured responses, but also that neural responses are not uniform in amplitude across eccentricity (Ales et al., 2013) or polar angle (Liu, Heeger, & Carrasco, 2006).

Accounting for these possibilities would likely lead to closer matches between data and model. Similarly, allowing the degree of synchrony to vary between 0 and 100% might lead to greater overlap in the predicted and observed spatial topography across MEG sensors.

Other factors which limit model accuracy of the current simulations include errors in MEG-MRI alignment and simplifications assumed in the head models. An important goal in future work will be to express simulation frameworks like our encoding model presented here with low-dimensional parameterizations that can be fit to data, and to optimize each stage of the computations to maximize model accuracy.

### 3.8 Conclusion

The ability to measure human brain activity at high temporal resolution has value to scientists and clinicians in many fields. EEG and MEG are important instruments for this reason. Typically, temporal properties of the EEG or MEG sensor responses are used to make inferences about the temporal dynamics of the underlying neural responses, and spatial properties of the sensor responses are used to infer spatial properties of the cortical responses. Our study showed that the *spatial* pattern of sensor responses can be used to make inferences about the *temporal* pattern of neural responses. We were able to make this observation through simulations with a visual encoding model, which showed that the degree of large-scale neural synchrony across cortex can have a large impact on the spatial topography of MEG sensor responses. This modeling result, combined with experimental data, allowed us to make inferences about the degree of large-scale synchrony underlying two types of visually driven MEG measures, stimulus-locked and broadband responses. We infer that the former is highly synchronized across cortex, whereas the latter is largely asynchronous.

The modeling approach we developed can be extended and applied to a wide range of paradigms to test specific and constrained hypothesis about the spatiotemporal pattern of neural responses measured non-invasively in the living human brain.

## 4. Methods

We re-analyzed and extended a published data set (Kupers et al., 2018). The extensions include adding anatomical MRI data for subjects who only had MEG data in the prior paper, and adding new subjects with both MRI and MEG data. The collection of new data adhered as closely as possible to the previously published methods. Several of the sections below - *Stimuli*, *Experimental design*, *MEG Data acquisition*, and *MEG preprocessing* – are highly similar to text in the previous paper since they describe the same experiment.

### 4.1 Subjects

The previous study included 8 subjects from New York University measured with MEG. We were able to recruit 6 of these subjects to participate in anatomical MRI measurements. We also recruited 6 new subjects for both MEG and MRI. The combined set of 12 subjects includes 8 females, ages 20-42 years (M = 28.3 / SD = 6.4 years) with normal or corrected-to-normal vision. All subjects provided written informed consent. The experimental protocol was in compliance with the safety guidelines for MRI and MEG research and was approved by the University Committee on Activities involving Human Subjects at New York University.

### 4.2 Stimuli

Stimuli were contrast-reversing dartboard patterns (12 square wave contrast-reversals per second), windowed within a circular aperture with a diameter of 22 degrees. (The previous study also included blocks with half-circle apertures, but these were not used for the new subjects and thus are not analyzed in this paper.) A uniform gray equal to the mean luminance of the black and white checks (206 cd/m²) was the background for the dartboards and was shown in the full screen during blank blocks. For more details, see methods and figure 1 of (Kupers et al., 2018).

### 4.3 Experimental design

Experiments consisted of multiple runs, 72 seconds each, containing alternating blocks of stimulation (contrast-reversing dartboards) and blanks (uniform gray field), 6 seconds each. There was a fixation dot in the middle of the screen throughout the run, switching between red and green at random intervals (averaging 3 seconds). The subjects were instructed to maintain fixation throughout the run and press a button every time the fixation dot changed color. The subjects were asked to minimize their blinking and head movements during the 72-s runs. After each run, there was a short break to blink and relax (typically 30-s to 1 minute).

In the previous study (Kupers et al., 2018), we obtained fifteen runs per subject. Within each run, the six stimulus blocks included two with full-circle apertures, two with left semicircular apertures, and two with right semicircular apertures. For the current study, only the full-circle apertures were analyzed. For the six new datasets, we obtained fewer runs (10 per subject rather than 15), but the left and right semicircular apertures were replaced with the full-circle apertures. In total, there were 30 full-circle stimulus blocks per subject for the prior datasets (15 runs x 2 stimulus blocks, counting only the stimulus blocks with full-circle apertures) and 60 for the new datasets (10 runs x 6 stimulus blocks).

### 4.4 Data acquisition

#### 4.4.1 MEG

The MEG data were acquired with a whole head Yokogawa MEG system (Kanazawa Institute of Technology, Japan) containing 157 axial gradiometers. The measurements were recorded at 1000 Hz with online high-pass and low-pass filters. The high-pass cutoff was 1 Hz, and the low-pass cutoff was either 200 Hz (prior study) or 500 Hz (new datasets). For registration of the head, we measured each subject’s head shape prior to the MEG scan using a handheld FastSCAN laser scanner (Polhemus, VT, USA). We used the laser scanner to digitize 8 locations: 3 on the forehead, the nasion, the left and right tragus, and the left and right peri-auricular points, and marked these locations with a non-permanent pen. We then placed 5 electrodes on the same locations on the forehead and peri-auricular points.

Before and after the main MEG experiment, recordings were made of the marker locations (electrodes) within the MEG dewar.

#### 4.4.2 MRI

Structural MRI scanning was conducted at the New York University Center for Brain Imaging using either a 3T Siemens Allegra (subjects S1, S3-S6), or a 3T Siemens Prisma (subjects S2, S7-S12). High resolution T1-weighted (T1w) whole brain anatomical images (1 mm^3^ isotropic voxels) were collected for each subject with a 3D rapid gradient echo (or ‘MPRAGE’) sequence.

### 4.5 Data analysis

#### 4.5.1 Reproducible computation and code sharing

All analyses were conducted in MATLAB (MathWorks, MA, USA). The analysis code and data are made publicly available via the Open Science Framework at the URL (https://osf.io/52mqt/). The code on this site includes scripts to reproduce all figures from the minimally pre-processed data. Each data figure has a single script named *makeFigureX* (where ‘*X’* is the figure number).

#### 4.5.2 MEG preprocessing

Raw MEG files were read into MATLAB by the FieldTrip Toolbox (Oostenveld, Fries, Maris, & Schoffelen, 2011 & Schoffelen, 2011). MEG data contained 6-s stimulus and 6-s blank blocks, which were sub-divided into 1-s non-overlapping epochs. The first 1-s epoch in each block was excluded from analysis to avoid the transient response caused by a change of the stimulus.

We then detected and removed outlier sensors and epochs by an automated routine This routine was also used in our previous study (Kupers et al., 2018) and is publicly available in the code repository linked on the OSF website (https://osf.io/52mqt/<u>, function</u> *nppPreprocessData.m*). This routine computes the variance of the time series within each 1-s epoch and labels as ‘bad’ those epochs having a variance that was either 20 times smaller or 20 times larger than the median variance across all epochs. If more than 20% of the epochs were ‘bad’ across individual sensors, the entire epoch was removed from the analysis. Similarly, if more than 20% of the epochs were labeled ‘bad’ within individual sensors, the entire sensor was removed. For those ‘bad’ epochs that were not removed by previous thresholds, the 1-s time series replaced by the spatially weighted interpolation of time series from nearby sensors (*i.e.*, weighting sensors inversely with the distance). On average, this automated procedure removed 4.9 sensors and 2.3% of the 1-s epochs per subject session due to noise or defect sensors. This preprocessing step is similar to other preprocessing algorithms (*e.g.* ‘Autoreject’ by (Jas et al., 2018)) and we believe that any such algorithm could substitute our preprocessing routine.

MEG data were separated into two data components: a stimulus-locked and broadband value. For comparison of the two signal types, we choose to analyze both types in units of amplitude. Although we defined the broadband response in terms of the power spectrum in prior work (Winawer et al., 2013; Kupers et al., 2018) (the square of the amplitude spectrum), we choose units of amplitude for both components in this paper. The amplitude domain is justified because both the stimulus-locked and broadband responses were normally distributed in amplitude, whereas only broadband (not stimulus-locked) distributions were normally distributed in power. Normally distributed variables are better suited for statistical summaries than skewed distributions. Nonetheless, the pattern of results does not change if we compute both data components as power, or stimulus-locked responses in amplitude and broadband responses in power.

For all but one subject (S10), the stimulus-locked component was computed as in the previous study. In short, we computed the Fast Fourier Transform of the time series within each 1-s epoch, extracted 12 Hz amplitudes for every epoch, and averaged this value across epochs separately for stimulus and blank. Data were then summarized into one value per sensor as the difference in amplitude between averaged stimulus and blank epochs.

For subject S10, we computed the stimulus-locked amplitudes *after* averaging the time series across epochs (sometimes called the coherent spectrum). This was because the MEG data for this subject contained a large alpha response close to the stimulus frequency, thereby masking the stimulus-locked response. Because the stimulus locked response has about the same phase in each epoch, but the alpha rhythm does not, averaging in time reduces the alpha rhythm much more than the 12-Hz stimulus locked response. After averaging in time and the computing the 12 Hz amplitude from the Fourier transformed time series, the stimulus-locked component was summarized as for the other subjects described above. If instead we remove S10 entirely from the analysis, or if we compute the coherent signal for all subjects, the spatial topography of the group average stimulus-locked signal is largely unchanged.

The computation for the broadband component was identical for all subjects and both studies: we took the geometric mean of the log amplitude within 60-150 Hz, excluding frequencies that could potentially include stimulus-locked harmonics (multiples of 12). As with the stimulus locked response, this resulted in one value per epoch per sensor, which were averaged separately for stimulus and blank periods and summarized as the difference between the stimulus and blank amplitudes.

To increase the signal-to-noise ratio of the broadband response, we applied a denoising algorithm on individual 1-s epochs developed in our lab previously (Kupers et al., 2018). This denoising algorithm, called NoisePool-PCA, identifies sensors that show small or no response to the stimulus as the ‘noise pool’. Data from the noise pool are used to compute noise components for each 1-s epoch using principal components analysis. These noise components are then projected out from all sensor time series for each 1-s epoch. For every subject, we projected out the number of PCs that gave the highest coefficient of determination, or the first 10 PCs if no maximum was reached within the first 10 PCs. The denoising algorithm increases the SNR in the broadband response, but does not cause a systematic change in the spatial pattern of the sensor responses. This is evident from figures 9 and 10 in Kupers et al. (2018), top row.

#### 4.5.3 MRI preprocessing

T1-weighted anatomy scans were co-registered and segmented into gray and white matter voxels using FreeSurfer’s *recon-all* auto-segmentation algorithm (Dale, Fischl, & Sereno, 1999; Fischl, Sereno, & Dale, 1999). For each individual subject’s cortical surface, we applied anatomical templates of retinotopy using the publicly available algorithm published by Benson *et al*. (2014). This algorithm uses an algebraic model of retinotopy on the flattened cortical surface anatomy, resulting in a template for occipital areas V1, V2, and V3, with their corresponding eccentricity and polar angle representation. The eccentricity template ranges from 0.1 to 80 degrees in visual angle and the polar angle template from 0 to 360 degrees. The Benson *et al*. algorithm is implemented in a Docker (https://hub.docker.com/r/nben/occipital_atlas) and does not need any manual intervention. The V1-V3 templates are reported to be at least as accurate as visual maps based off 6.4 minutes of retinotopy scans (Benson & Winawer, 2018).

To check the contribution of higher visual areas to the spatial topography of predicted responses in MEG sensors, we ran our model simulations with two more atlases (Supplementary Figure S7). The first additional atlas was the extended version of the Benson *et al*. templates (Benson & Winawer, 2018) providing polar angle and eccentricity maps for V1, V2, V3, hV4, VO1/2, LO1/2, TO1/2, V3A/B. This allowed us to limit the cortical sources to the stimulus aperture in every visual area (0.18-11 degrees). The second atlas was a probabilistic atlas of visual areas provided by Wang and colleagues (Wang, Mruczek, Arcaro, & Kastner, 2015). This atlas provides the borders of 25 visual areas: V1v/d, V2v/d, V3v/d, hV4, VO1/2, PHC1/2, TO1/2, LO1/2, V3A/B, IPS0-5, SPL1, and FEF. These areas were defined by an experiment with a stimulus extent of ∼15 degrees eccentricity.

#### 4.5.4 MRI-MEG alignment

We used the Brainstorm toolbox (Tadel et al., 2011) to align the T1-weighted anatomy to the MEG helmet and the head using 6 fiducials points: the nasion, left/right peri-auricular, interhemispheric, anterior and posterior commissure. This procedure resulted in a single coordinate space for all three measurements.

When importing subject’s FreeSurfer folder into the Brainstorm toolbox, the T1w anatomy and cortical surfaces are by default downsampled to 7,501 vertices per hemisphere. We likewise downsampled the retinotopic atlases to the same resolution using nearest-neighbor interpolation.

#### 4.5.5 V1-V3 source time series simulation using noiseless sine waves

For simulations in Figure 5 and 8, the cortical time series for responsive voxels (*i.e.*, in the V1-V3 maps and with population receptive field (pRF) centers within the stimulus aperture) were sine waves of unit amplitude and frequency of one cycle per epoch. The time series for all other vertices was fixed at 0. We generated 1,000 epochs for a simulation, each consisting of 10 time points.

For the synchronous simulations, the phase of each sine wave was identical. For asynchronous simulations, the phase was randomized across epochs. For Supplementary Figure S9, we simulated time series with intermediate levels of synchrony. Here, the phases were sampled from Von Mises distributions varying in width (kappa parameter) between all asynchronous (kappa=0) and all synchronous (kappa=100*pi).

#### 4.5.6 V1-V3 source time series simulation using a more complete, biologically plausible generation of cortical responses

In addition to simulating cortical time series using noiseless sine waves, we also implemented a more detailed simulation of synchronous and asynchronous V1-V3 responses based on neural biophysics (**Supplementary Figure S6**). This simulation generates ECoG responses in visual cortex to full-field contrast-reversing dartboard patterns and is adapted from the simulation previously published by Winawer *et al*. (2013). In brief, we first simulate incoming spikes generated by a Poisson process. Each spike elicits a synaptic current assuming a time-invariant linear system, characterized by a gamma impulse response function (exponent 2, time constant 0.0023 s). These currents are then summed by the dendrite as a leaky integrator (time constant alpha = 0.1 s). The time constants are identical to those by Winawer *et al*. (2013) and based on a specific version of the ECoG simulation published by Miller *et al*. (Miller et al., 2009).

The time-varying rate of the spike arrivals is modeled as the sum of three process: an evoked response, an induced response, and spontaneous activity. The evoked response rate is an impulse response function convolved with the stimulus reversal events. The induced response is an elevated rate whenever the stimulus is present. The induced activity is constant across time. The rates are 15 spikes/second/synapse for the evoked and induced response, and 10 spikes/second/synapse for the spontaneous activity. (For the evoked response, the rate is the mean across time). We simulated 100 1-s epochs with 1-ms time bins. Just as in our simple simulation, we simulate these ECoG responses for each vertex within V1-V3 and multiply the responses with individual subject’s gain matrix to get predicted MEG sensor responses (see next section 4.5.7). The exact details of all steps in simulation are in the MATLAB script, *makeSupplementaryFigure6.m*.

To analyze the simulated data, we extract the stimulus-locked and broadband amplitudes from stimulus and blank periods in MEG sensors, using the same procedure as for the analysis of actual MEG data with one modification. For the stimulus-locked response, we subtracted the estimated broadband component. This is because the simulated broadband response, unlike the observed MEG broadband response, included temporal frequencies that overlapped the stimulus frequency.

#### 4.5.7 Forward model (sources to sensors)

After the alignment of the T1w anatomy and MEG sensor space, we computed the gain matrix (also known as ‘head model’ or ‘lead field matrix’) with Brainstorm’s MEG Overlapping Spheres method (Huang et al., 1999). This algorithm fits one local sphere under each MEG sensor, resulting in a weighted sum of cortical locations in FreeSurfer’s downsampled pial surface contributing to that MEG sensor. For panel A and B in **Supplementary Figure S7**, we use the three-shells Boundary Element Model (BEM) instead (Kybic et al., 2005; Gramfort et al., 2010). The constrained gain matrix derived from BEM contained the same number of vertices as the Overlapping Spheres head model.

For both head models, the gain matrices were limited to one orientation per vertex perpendicular to the cortical surface.

To predict the responses of the MEG sensors, we use the following equation:

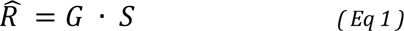

where *R̂* is the predicted sensor responses (*k* time points x *m* sensors), in this case 10,000 time points (1,000 epochs of 10 time points) by 157 sensors. *G* is the constrained gain matrix computed by Brainstorm (*n* sources x *m* sensors), in this case 15,002 sources by 157 sensors. *S* is the source activity (*k* time points by *n* sources), in this case 10,000 time points (1,000 epochs of 10 time points each) by 15,002 sources.

#### 4.5.8 Summary metrics and visualization

To summarize the simulated neural responses predicted by the forward, we computed the amplitude at the input frequency of the sine wave (1 cycle per epoch) using the Fourier transform of the predicted responses *R̂* for the V1-V3 time series. This was done separately for synchronous or asynchronous simulated time series. We averaged the amplitudes across epochs for each sensor as for MEG data. MEG data were summarized in individual subjects into stimulus-locked and broadband amplitudes. For every subject, we subtract the mean signal across blank epochs from the mean signal across stimulus epochs.

Because the stimulus-locked frequency (12 Hz) is within the typical alpha range, one might be concerned that our contrast summary measure reflects the endogenous alpha rhythm and not the stimulus-driven response. To check for this, we compared the responses at the stimulus frequency and a nearby frequency (12 Hz and 11 Hz), both when the stimulus is present and when it is absent (blanks). We find that when the stimulus is present, the amplitude at the stimulus frequency is about 20 times larger than the amplitude at adjacent frequencies. During the blank, this is not the case - the amplitude is about the same across nearby frequencies in the alpha band. This is consistent with the 12 Hz amplitude during visual stimulation being largely driven by the stimulus. See *s_reviewFigures.m* for details.

The stimulus-locked and broadband amplitudes were thresholded by a signal-to-noise ratio (SNR) of 1. The SNR was calculated as the ratio of the mean to the standard deviation of the stimulus-locked or broadband summary measure across 1000 bootstraps. Group averages were calculated as the mean SNR across subjects.

We visualized the predictions or observed data for each sensor on a topographic sensor map using the FieldTrip toolbox (Oostenveld et al., 2011). Observed data have a color map restricted to the 97.5th percentile of the plotted response, predicted responses are normalized by the maximum of the synchronous sensor-wise average across subjects. We further plotted isocontour lines around the 10 sensors with the largest response (93.6^th^ percentile of the data or model predictions.

Topographic sensor maps were summarized as 1-dimensional line plots. A narrow, vertical Gaussian pooling window was used to compute a weighted average across all sensors for each horizontal bin. This pooling window averaged amplitudes for each sensor falling within the bin, using 100 equally sized bins from the most left to most right located MEG sensor. For individual observers, the average sensor amplitudes were computed from the contrast map (difference between average stimulus and average blank epochs for either stimulus-locked or broadband responses). Error bars were computed by bootstrapping the weighted average and taking the standard deviation across 1000 bootstraps for each bin. For the group average, we used the arithmetic mean across observers for stimulus-locked or broadband responses to compute the weighted group average and error bars representing the standard error across observers for each bin.

## 5. Supplementary Material

**Supplementary Figure S1.**
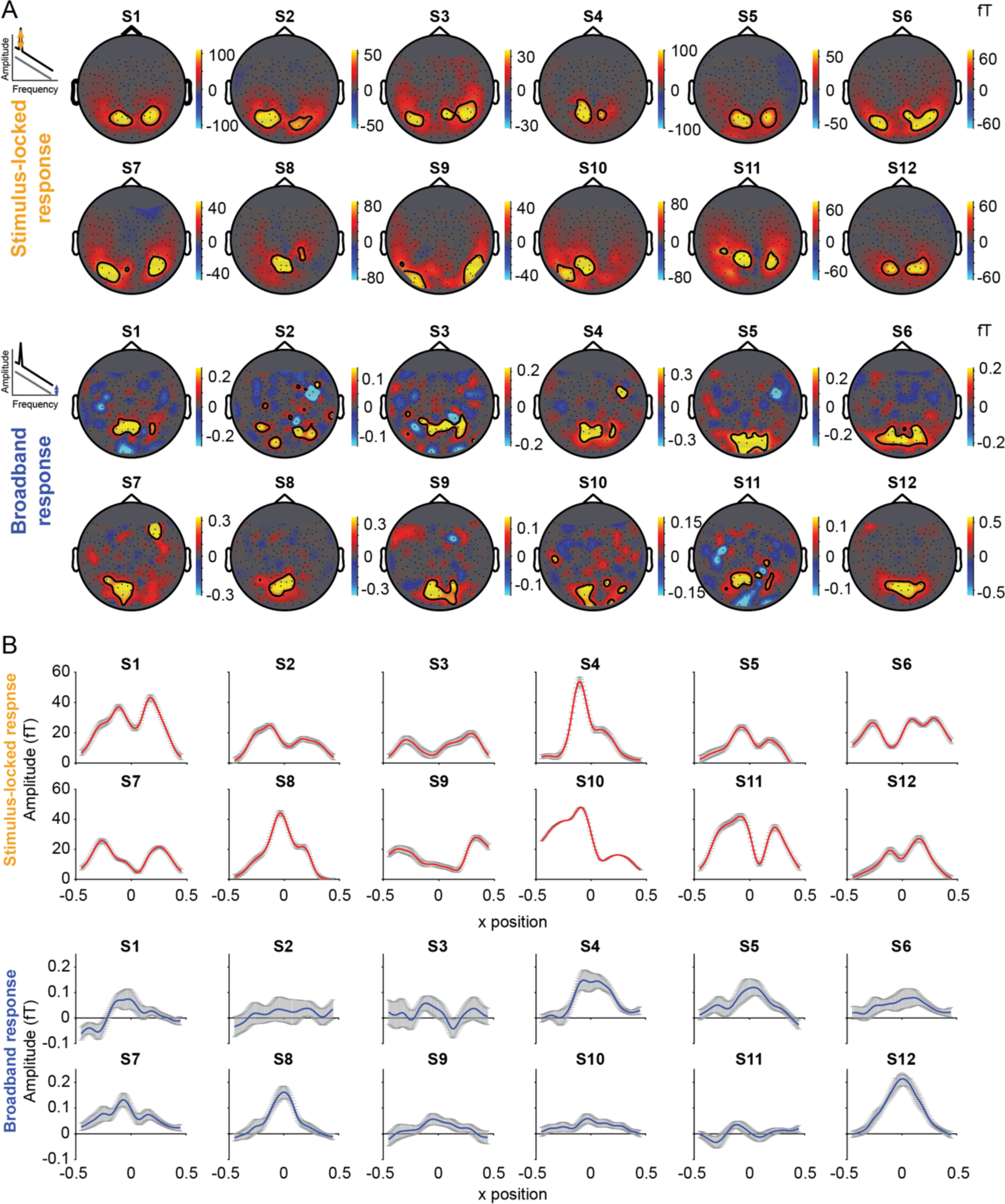
Observed stimulus-locked and broadband responses for all individual subjects. **(A)** Topographic MEG sensor maps for stimulus-locked (row 1-2) and broadband response (row 3-4). Both stimulus-locked and broadband responses are in femto Tesla amplitudes. **(B)** Weighted average of stimulus-locked (row 1-2) and broadband (row 3-4) response across posterior sensors. Colored line shows sample mean of moving window taking the Gaussian weighted average across posterior MEG sensors, error bars are standard deviation across bootstraps.

**Supplementary Figure S2.**
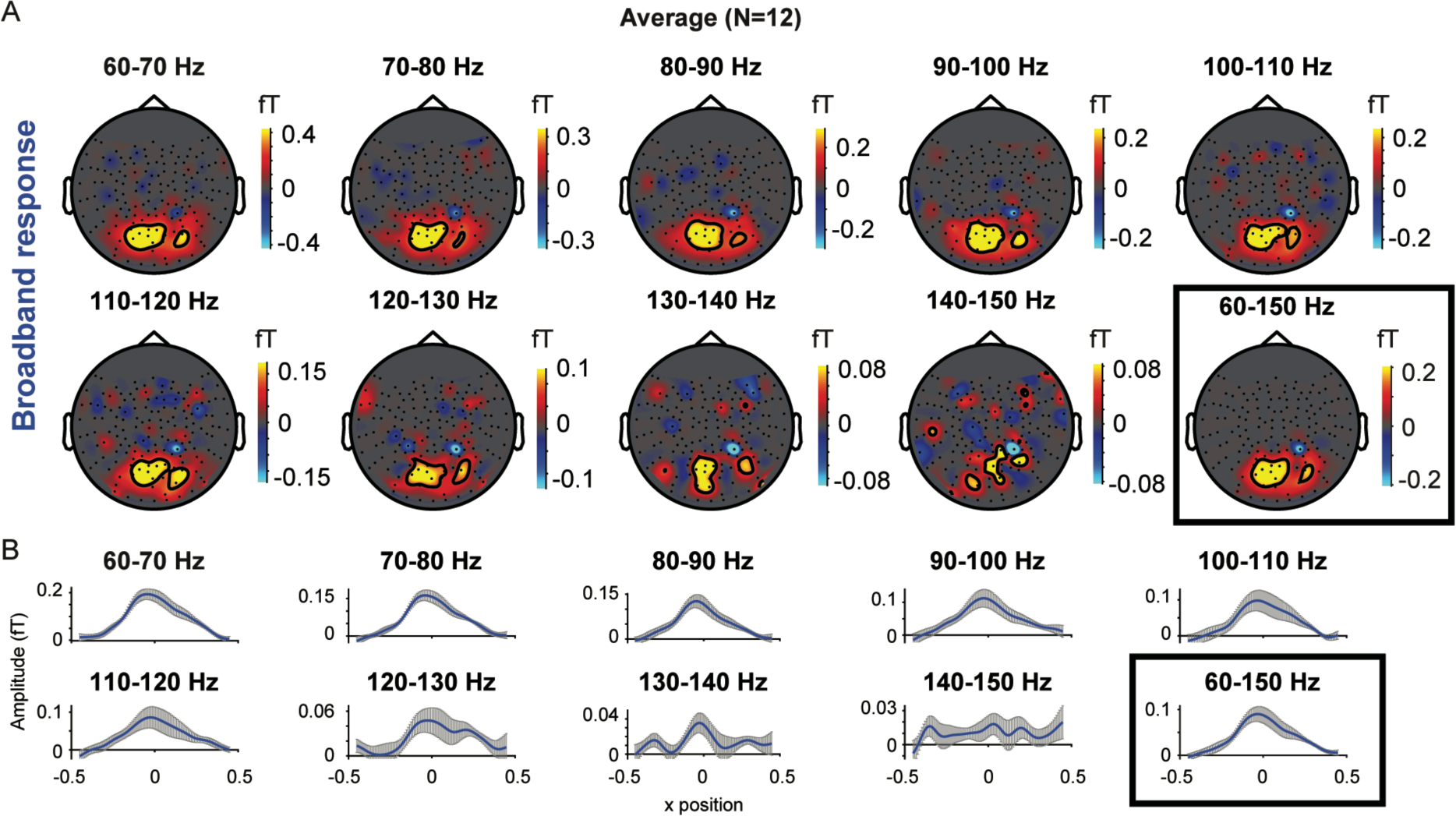
Group average broadband responses analyzed in separate, narrow frequency bands. **(A)** Topographic MEG sensor maps of broadband responses limited to 10-Hz frequency bands from 60-150 Hz, excluding stimulus harmonics. **(B)** Weighted average of broadband responses across sensors. The shaded line shows the sample mean and standard deviation across bootstraps. The black boxes highlight the frequency range used for analysis in the manuscript, 60-150 Hz. The spatial pattern in most of the narrow bins resembles the pattern in the wide bin.

**Supplementary Figure S3.**
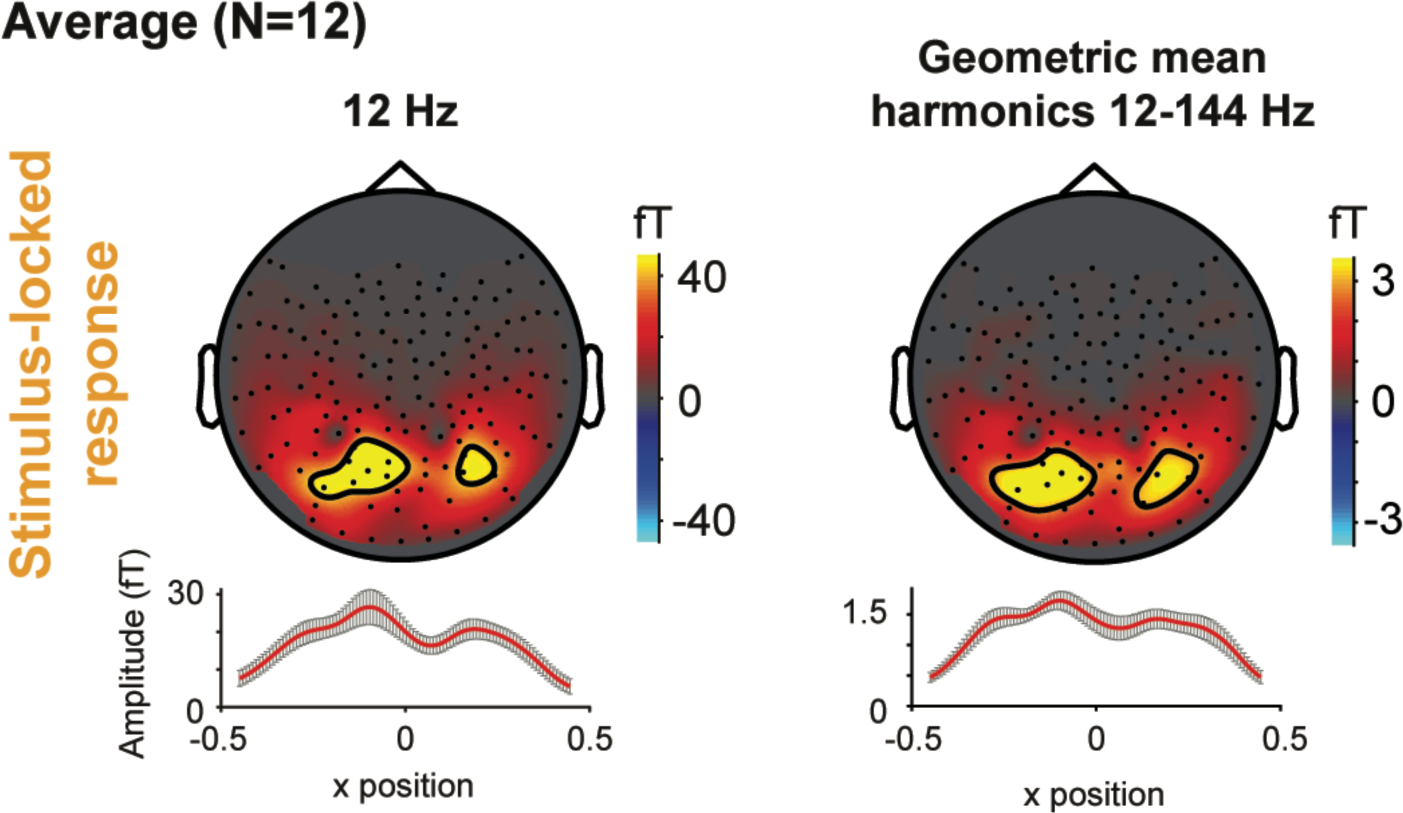
Group average stimulus-locked response for 12 Hz only versus all stimulus harmonics up to 144 Hz. **Left**: Identical to group average in Figure 3. Topographic MEG sensor maps for stimulus-locked response analyzed as the difference between stimulus and blank epochs at 12 Hz, with a 1D summary of the sensor responses shown below. **Right:** Same as left panel, but now the stimulus-locked response is computed as the geometric mean across all 12 Hz harmonics from 12 to 144 Hz (excluding 60 and 120 Hz as those frequencies are contaminated by line noise). Both the left and right panel show bimodal spatial distributions.

**Supplementary Figure S4.**
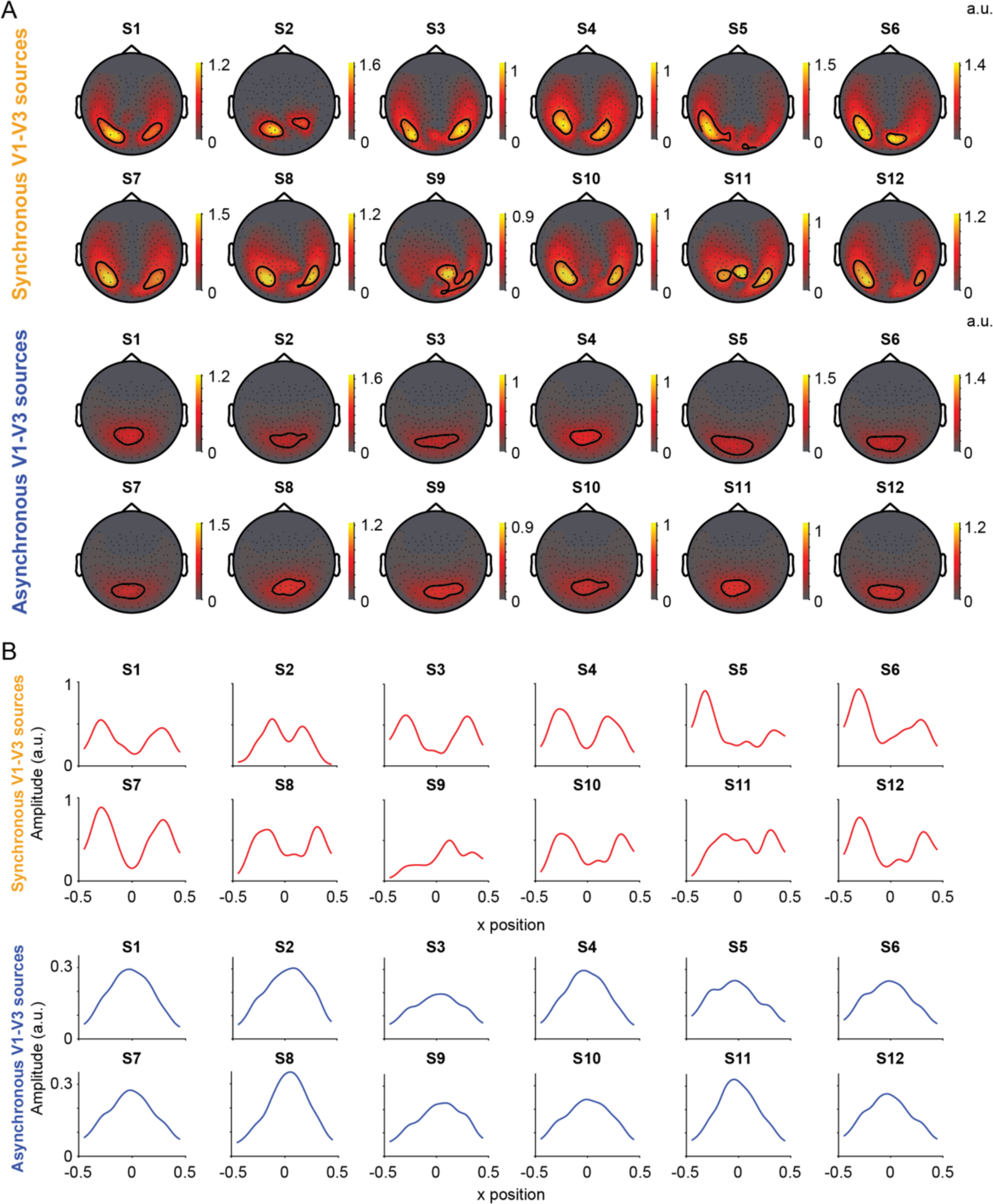
Forward model predictions of signals from V1-V3 to MEG sensors for all individual subjects. **(A)** Top panel: Model predictions using synchronous V1-V3 sources, all individual amplitudes are arbitrary units with the same scale factor shown in the upper right. Bottom panel: Same as top panel, but for asynchronous V1-V3 sources. All amplitudes are arbitrary units. Predicted sensor responses for both synchronous and asynchronous simulations are normalized to the largest response in the synchronous spatial map of the sensor-wise group average (see for example Figure 5A, top left panel). **(B)** 1D summary of the sensor responses in A.

**Supplementary Figure S5.**
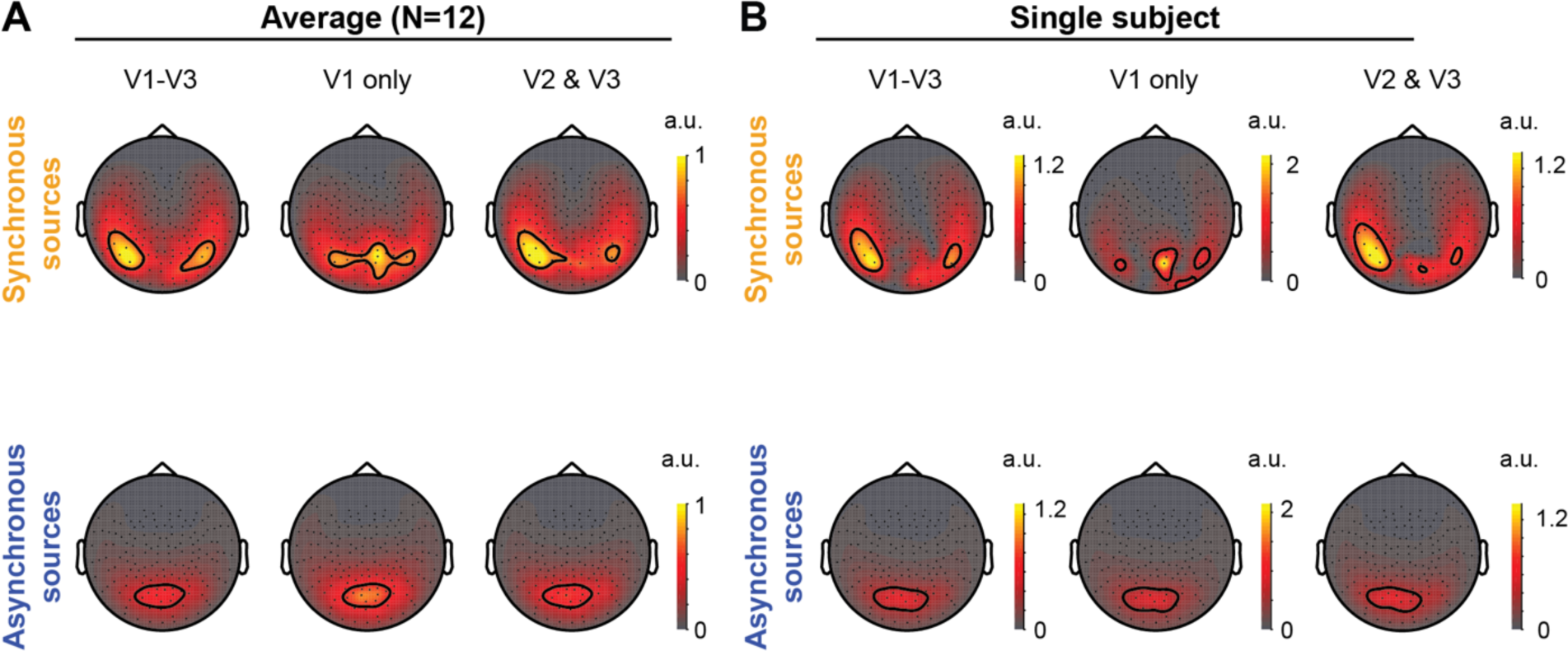
Forward model predictions of signals in MEG sensors comparing sources fromV1-V3, V1 only, and V2 and V3 without V1. Layout as in Figure 5. **(A)** Model predictions for average across 12 subjects using sensor-wise averaging. **(B)** Model predictions for a single subject (S12). All amplitudes are arbitrary units. Predicted sensor responses for both synchronous and asynchronous simulations are normalized to the largest response in the synchronous spatial map of the sensor-wise group average (top left panel in A).

**Supplementary Figure S6.**
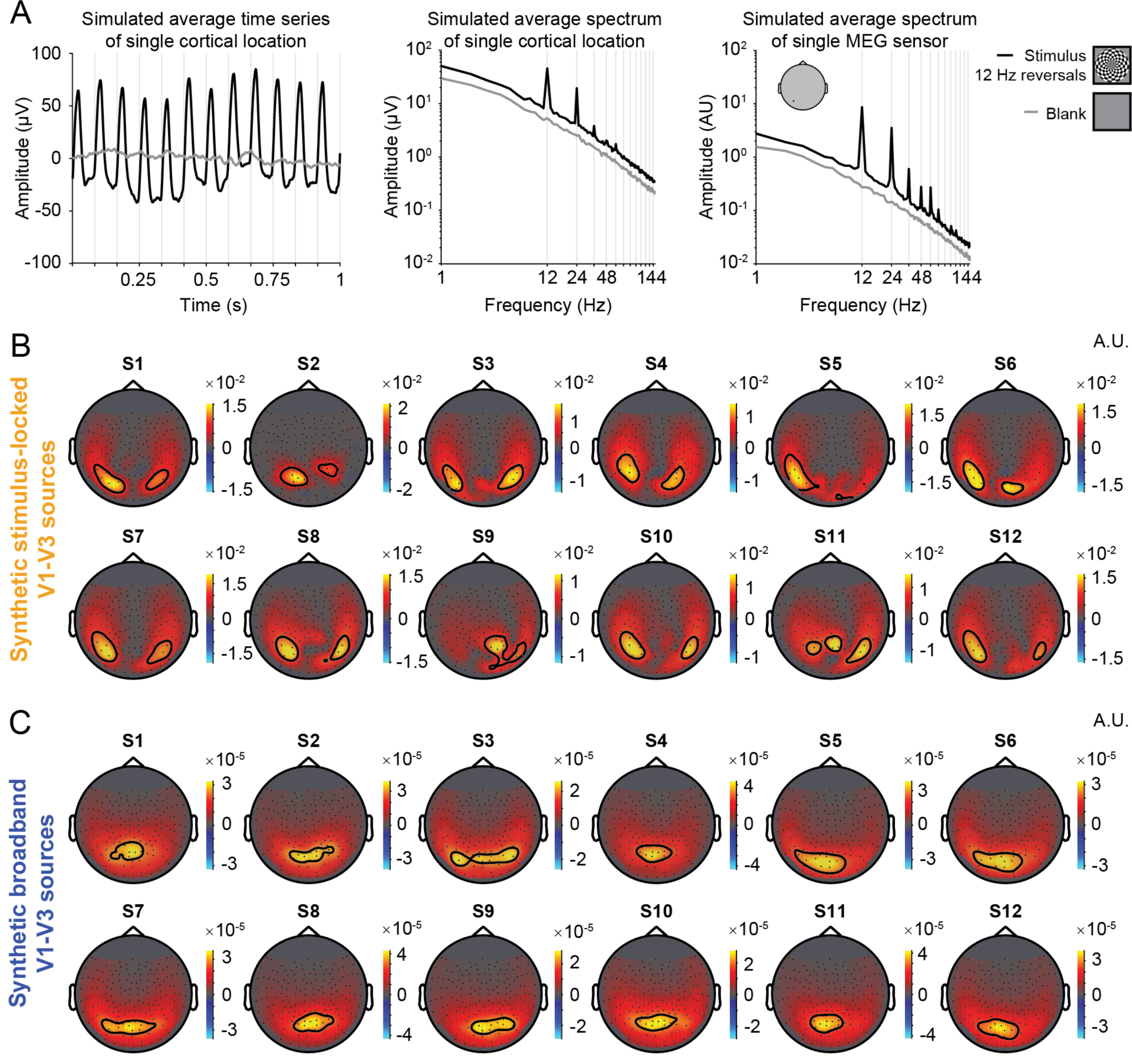
Forward model predictions for biologically plausible simulated cortical signals. We implemented a more detailed simulation of cortical timeseries adapted from previously published ECoG simulations by Winawer *et al*. (2013). **(A)** We generate cortical time series to 12 Hz contrast reversing dartboard patterns or no stimuli (blanks). **Left panel** shows the average time course of a single V1 vertex across 100 simulated stimulus epochs (black line) or blank epochs (gray line). **Middle panel** shows Fourier amplitudes averaged across stimulus or blank epochs of single cortical location. Simulated stimulus epochs contain clear stimulus-locked responses at 12 Hz and harmonics and a broadband response up to lower frequencies. Both stimulus-locked and broadband response fall off with a 1/f slope. The cortical time series are multiplied with the gain matrix from the Overlapping Spheres head model (Huang et al., 1999), resulting in the predicted MEG sensor time series. **Right panel** shows the Fourier amplitudes of a single MEG sensor. **(B)** Topographic maps showing the forward model predictions for the stimulus-locked component extracted from simulated ECoG signals in V1-V3 sources. **(C)** Same as panel B, but for the broadband component extracted from simulated ECoG signals in V1-V3 sources. Both panels show amplitudes in arbitrary units.

**Supplementary Figure S7.**
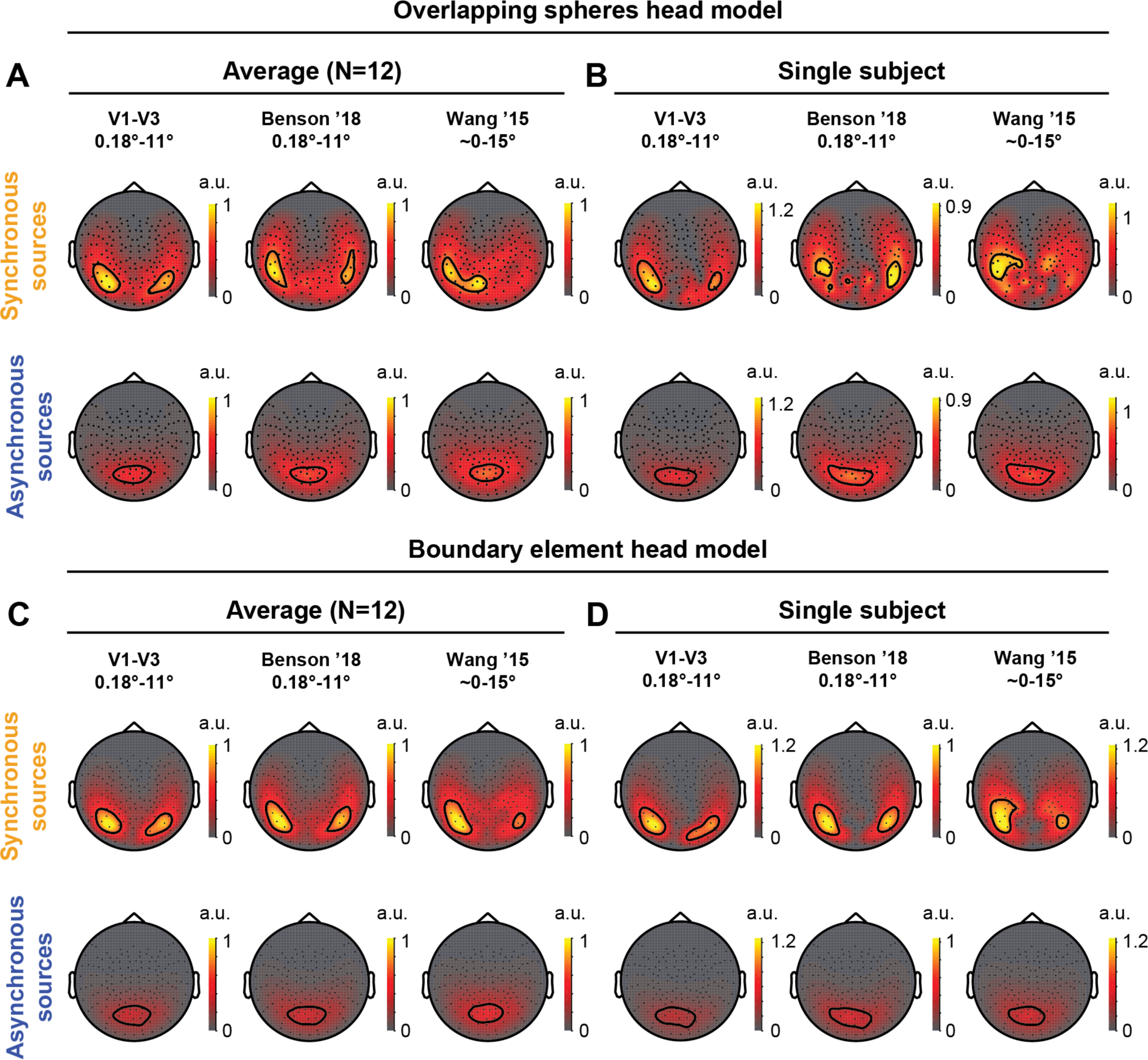
The effect of volume conduction head model and included ROIs on forward model predictions of signals in MEG sensors. **(A)** Forward model predictions for the average across 12 subjects using the overlapping spheres head model. This means that synchronous and asynchronous source activity were simulated and projected to the MEG sensors using a gain matrix computed from the overlapping spheres head model (Huang et al., 1999). Columns compare synchronous and asynchronous sources from V1-V3 (first column) or all 12 ROIs in the latest version of the Benson atlas (Benson & Winawer, 2018) (second column) or 25 ROIs in the probabilistic Wang *et al*. (2015) atlas (third column). Since the Wang *et al*. (2015) atlas only provides the outlines of visual areas, all vertices within the ROIs were used, corresponding to approximately 0 to 15 deg. **(B)** Same as in (A) but for single subject example. Data are from subject S12. All amplitudes are arbitrary units. **(C & D)** Similar to panel A and B, but using forward model predictions for the average across 12 subjects using the 3-layer boundary element model (Kybic et al., 2005; Gramfort et al., 2010). Group average topographies are averaged in sensor-space. Separate for each type of head model and ROI selection, predicted sensor responses for both synchronous and asynchronous simulations are normalized to the largest response in the synchronous spatial map of the sensor-wise group average (top panels in A and C).

**Supplementary Figure S8.**
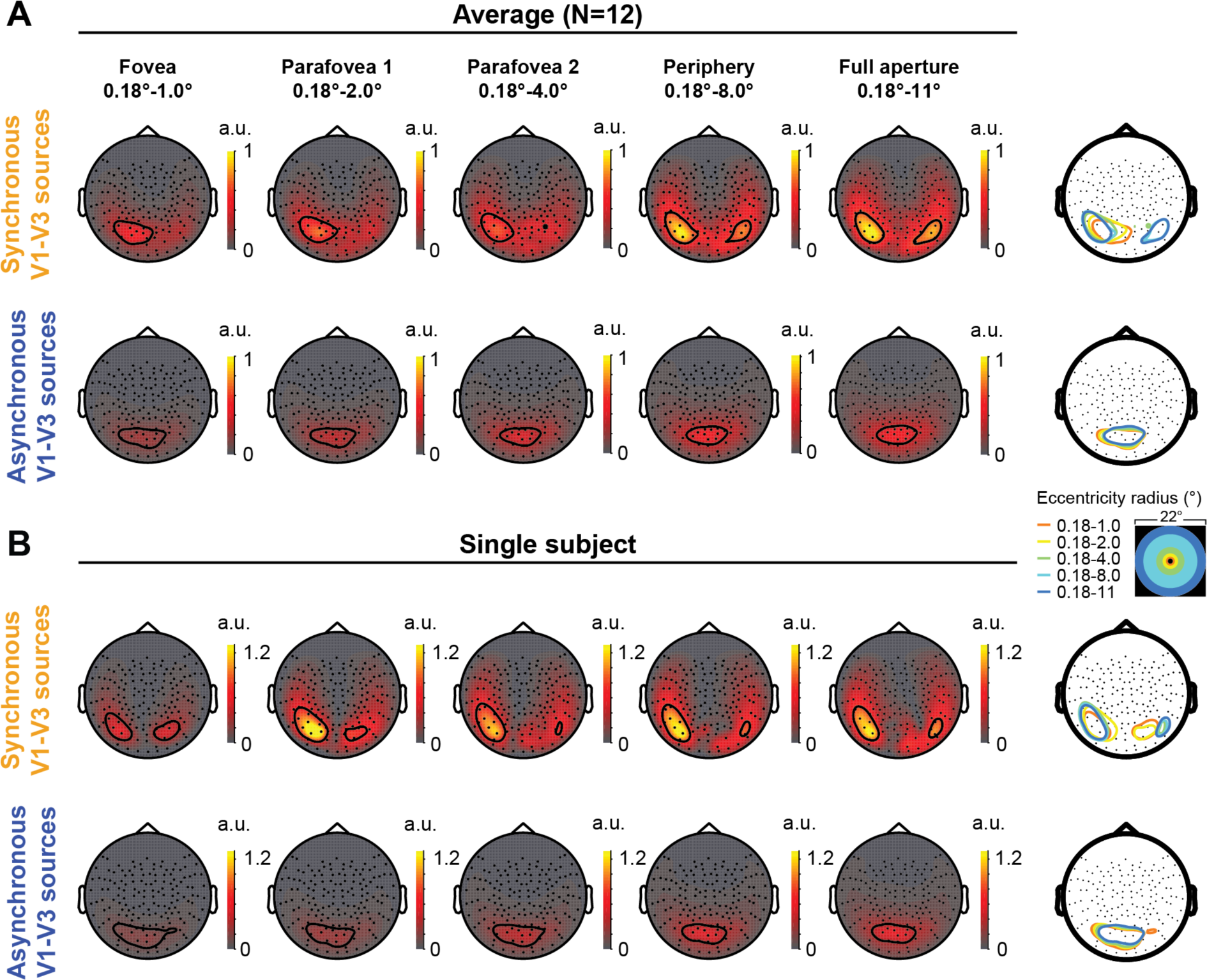
The effect of eccentricity on forward model predictions of signals in MEG sensors. **(A)** Forward model predictions for synchronous (top) and asynchronous (bottom) V1-V3 sources, averaged across 12 subjects. Columns represent predictions for different eccentricity bands accumulating in width from a foveal ring (left) to the entire stimulus aperture (right). Last column replots the contour lines for all eccentricity bands in a single mesh. Data are averaged in sensor-space. **(B)** Same as in (A) but for a single subject example, data are from subject S12. All amplitudes are arbitrary units. Predicted sensor responses for both synchronous and asynchronous simulations are normalized to the largest response in the synchronous spatial map of the sensor-wise group average (“Full aperture”, top right panel in A).

**Supplementary Figure S9.**
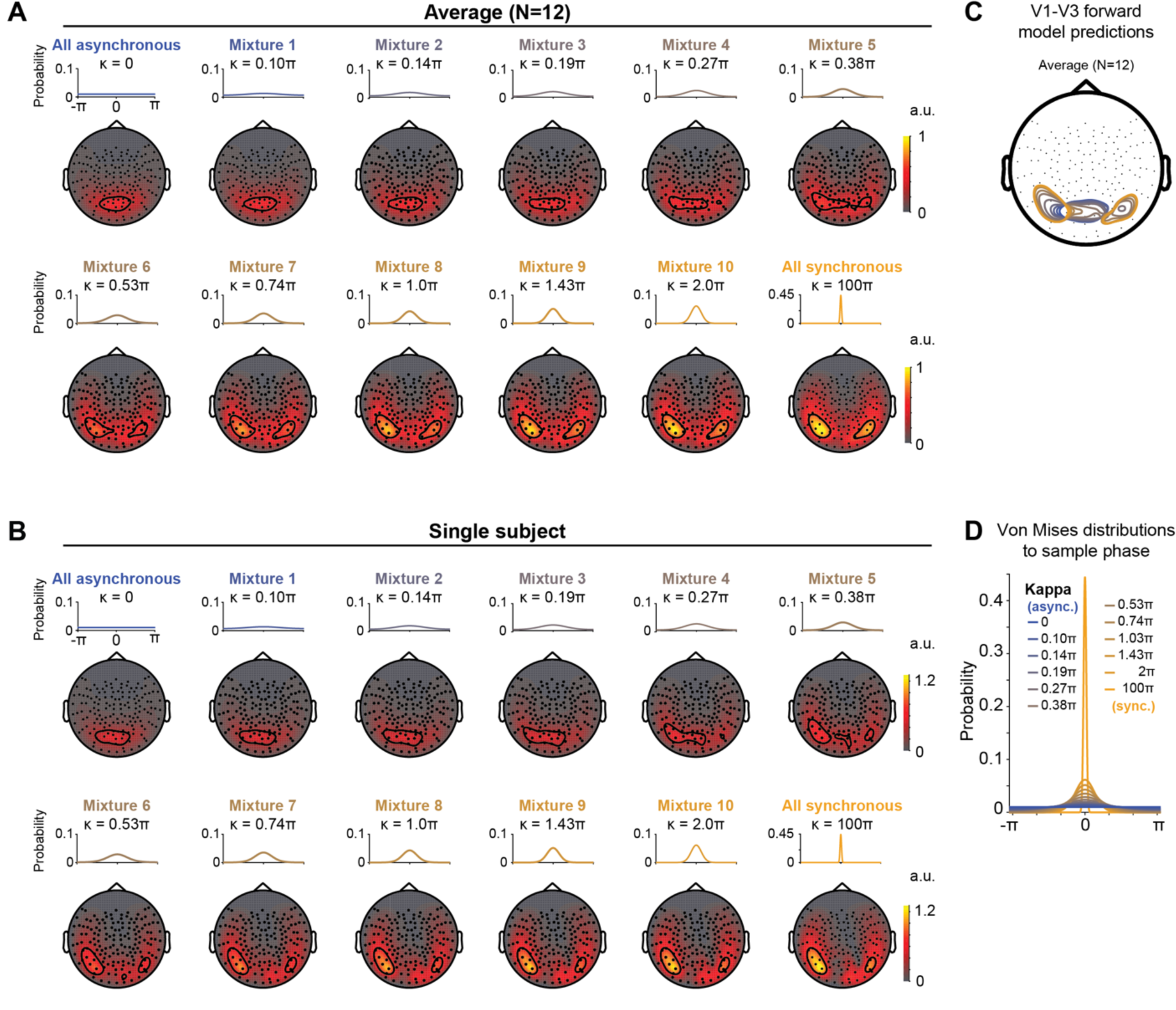
The effect of intermediate source synchrony levels on forward model predictions of signals in MEG sensors. **(A)** Forward model predictions for the average across 12 subjects. From left to right, rows show model predictions for source activity where its phase was sampled from a Von Mises distribution that decreases in width. A Von Mises distribution with a kappa of 0 gives a uniform distribution between –pi and pi, resulting in asynchronous source activity with randomized phases. The narrowest Von Mises distribution (kappa = 100*pi) has a FWHM of 0.13*pi and was used to sample phases for synchronous sources. Data are averaged in sensor-space. **(B)** Same as (A) but for a single subject. Data are from subject S12. All amplitudes are arbitrary units and normalized to the largest sensor response in the fully (100*pi) synchronous sensor-wise group average (“All synchronous”, lower right panel in A). **(C)** Replotted contour lines from forward model predictions using V1-V3 sources, averaged across 12 subjects, for the 12 Von Mises distributions. Colors represent the different Von Mises distributions used in forward model predictions as shown in panel D below. **(D)** Von Mises distributions to sample the phases of asynchronous (kappa=0, blue), synchronous (kappa=100*pi, orange) or 10 mixtures between the two extremes. Kappa values for mixtures were log spaced between 0 and 100*pi.

## References

Acar, Z. A., Makeig, S., & Worrell, G. (2008). Head modeling and cortical source localization in epilepsy. Conf Proc IEEE Eng Med Biol Soc, 2008, 3763–3766. doi: 10.1109/IEMBS.2008.4650027

Adrian, E. D., & Matthews, B. H. C. (1934). The Berger rhythm potential changes from the occipital lobes in man. Brain, 57, 355–385. doi: 10.1093/brain/57.4.355

Ahlfors, S. P., Han, J., Lin, F. H., Witzel, T., Belliveau, J. W., Hamalainen, M. S., & Halgren, E. (2010). Cancellation of EEG and MEG signals generated by extended and distributed sources. Human brain mapping, 31(1), 140–149. doi: 10.1002/hbm.20851

Akalin Acar, Z., & Makeig, S. (2013). Effects of forward model errors on EEG source localization. Brain Topogr, 26(3), 378–396. doi: 10.1007/s10548-012-0274-6

Ales, J. M., Carney, T., & Klein, S. A. (2010). The folding fingerprint of visual cortex reveals the timing of human V1 and V2. Neuroimage, 49(3), 2494–2502. doi: 10.1016/j.neuroimage.2009.09.022

Ales, J. M., Farzin, F., Rossion, B., & Norcia, A. M. (2012). An objective method for measuring face detection thresholds using the sweep steady-state visual evoked response. J Vis, 12(10). doi: 10.1167/12.10.18

Ales, J. M., Yates, J. L., & Norcia, A. M. (2010). V1 is not uniquely identified by polarity reversals of responses to upper and lower visual field stimuli. Neuroimage, 52(4), 1401–1409. doi: 10.1016/j.neuroimage.2010.05.016

Ales, J. M., Yates, J. L., & Norcia, A. M. (2013). On determining the intracranial sources of visual evoked potentials from scalp topography: a reply to Kelly et al. (this issue). Neuroimage, 64, 703–711. doi: 10.1016/j.neuroimage.2012.09.009

Bartoli, E., Bosking, W., Chen, Y., Li, Y., Sheth, S. A., Beauchamp, M. S., … Foster, B. L. (2019). Functionally Distinct Gamma Range Activity Revealed by Stimulus Tuning in Human Visual Cortex. Curr Biol, 29(20), 3345–3358 e3347. doi: 10.1016/j.cub.2019.08.004

Barzegaran, E., Bosse, S., & Norcia, A. M. (2019). EEGSourceSim: A framework for realistic simulation of EEG scalp data using MRI-based forward models and biologically plausible signals and noise. J Neurosci Methods, 108377. doi: 10.1016/j.jneumeth.2019.108377

Benson, N. C., Broderick, W. F., Müller, H., & Winawer, J. (2017). Toward a standard cortical observer. Paper presented at the OSA Fall Vision Meeting, Washington, D.C.

Benson, N. C., Butt, O. H., Brainard, D. H., & Aguirre, G. K. (2014). Correction of distortion in flattened representations of the cortical surface allows prediction of V1-V3 functional organization from anatomy. PLoS Comput Biol, 10(3), e1003538. doi: 10.1371/journal.pcbi.1003538

Benson, N. C., & Winawer, J. (2018). Bayesian analysis of retinotopic maps. Elife, 7. doi: 10.7554/eLife.40224

Berger, H. (1929). Über Elektroenkephalogramm des Menschen. European Archives of Psychiatry and Clinical Neuroscience, 87(1), 527–570. doi: 10.1007/BF01835097

Birbaumer, N., Elbert, T., Canavan, A. G., & Rockstroh, B. (1990). Slow potentials of the cerebral cortex and behavior. Physiol Rev, 70(1), 1–41. doi: 10.1152/physrev.1990.70.1.1

Brookes, M. J., Zumer, J. M., Stevenson, C. M., Hale, J. R., Barnes, G. R., Vrba, J., & Morris, P. G. (2010). Investigating spatial specificity and data averaging in MEG. Neuroimage, 49(1), 525–538. doi: 10.1016/j.neuroimage.2009.07.043

Butler, R., Bernier, P. M., Lefebvre, J., Gilbert, G., & Whittingstall, K. (2017). Decorrelated Input Dissociates Narrow Band gamma Power and BOLD in Human Visual Cortex. J Neurosci, 37(22), 5408–5418. doi: 10.1523/JNEUROSCI.3938-16.2017

Butler, R., Bernier, P. M., Mierzwinski, G. W., Descoteaux, M., Gilbert, G., & Whittingstall, K. (2019). Cortical distance, not cancellation, dominates inter-subject EEG gamma rhythm amplitude. Neuroimage, 192, 156–165. doi: 10.1016/j.neuroimage.2019.03.010

Buzsaki, G., Anastassiou, C. A., & Koch, C. (2012). The origin of extracellular fields and currents--EEG, ECoG, LFP and spikes. Nat Rev Neurosci, 13(6), 407–420. doi: 10.1038/nrn3241

Carl, C., Acik, A., Konig, P., Engel, A. K., & Hipp, J. F. (2012). The saccadic spike artifact in MEG. Neuroimage, 59(2), 1657–1667. doi: 10.1016/j.neuroimage.2011.09.020

Chen, C. M., Lakatos, P., Shah, A. S., Mehta, A. D., Givre, S. J., Javitt, D. C., & Schroeder, C. E. (2007). Functional anatomy and interaction of fast and slow visual pathways in macaque monkeys. Cereb Cortex, 17(7), 1561–1569. doi: 10.1093/cercor/bhl067

Cichy, R. M., Khosla, A., Pantazis, D., Torralba, A., & Oliva, A. (2016). Comparison of deep neural networks to spatio-temporal cortical dynamics of human visual object recognition reveals hierarchical correspondence. Sci Rep, 6, 27755. doi: 10.1038/srep27755

Cichy, R. M., Pantazis, D., & Oliva, A. (2016). Similarity-Based Fusion of MEG and fMRI Reveals Spatio-Temporal Dynamics in Human Cortex During Visual Object Recognition. Cereb Cortex, 26(8), 3563–3579. doi: 10.1093/cercor/bhw135

Cicmil, N., Bridge, H., Parker, A. J., Woolrich, M. W., & Krug, K. (2014). Localization of MEG human brain responses to retinotopic visual stimuli with contrasting source reconstruction approaches. Front Neurosci, 8, 127. doi: 10.3389/fnins.2014.00127

Cottereau, B., Ales, J. M., & Norcia, A. M. (2015). How to use fMRI functional localizers to improve EEG/MEG source estimation. J Neurosci Methods, 250, 64–73. doi: 10.1016/j.jneumeth.2014.07.015

Cottereau, B., Lorenceau, J., Gramfort, A., Clerc, M., Thirion, B., & Baillet, S. (2011). Phase delays within visual cortex shape the response to steady-state visual stimulation. Neuroimage, 54(3), 1919–1929. doi: 10.1016/j.neuroimage.2010.10.004

Crone, N. E., Miglioretti, D. L., Gordon, B., & Lesser, R. P. (1998). Functional mapping of human sensorimotor cortex with electrocorticographic spectral analysis. II. Event-related synchronization in the gamma band. Brain, 121 ( *Pt 12*), 2301–2315. doi: 10.1093/brain/121.12.2301

Dale, A. M., Fischl, B., & Sereno, M. I. (1999). Cortical surface-based analysis. I. Segmentation and surface reconstruction. Neuroimage, 9(2), 179–194. doi: 10.1006/nimg.1998.0395

Di Russo, F., Pitzalis, S., Aprile, T., Spitoni, G., Patria, F., Stella, A., … Hillyard, S. A. (2007). Spatiotemporal analysis of the cortical sources of the steady-state visual evoked potential. Human brain mapping, 28(4), 323–334. doi: 10.1002/hbm.20276

Di Russo, F., Pitzalis, S., Spitoni, G., Aprile, T., Patria, F., Spinelli, D., & Hillyard, S. A. (2005). Identification of the neural sources of the pattern-reversal VEP. Neuroimage, 24(3), 874–886. doi: 10.1016/j.neuroimage.2004.09.029

Dumoulin, S. O., & Wandell, B. A. (2008). Population receptive field estimates in human visual cortex. Neuroimage, 39(2), 647–660. doi: 10.1016/j.neuroimage.2007.09.034

Duvernoy, H. M. (1999). The Human Brain: Surface, Three-Dimensional Sectional Anatomy with MRI, and Blood Supply. Austria: Springer Vienna.

Enroth-Cugell, C., & Robson, J. G. (1966). The contrast sensitivity of retinal ganglion cells of the cat. J Physiol, 187(3), 517–552. doi: 10.1113/jphysiol.1966.sp008107

Fischl, B., Sereno, M. I., & Dale, A. M. (1999). Cortical surface-based analysis. II: Inflation, flattening, and a surface-based coordinate system. Neuroimage, 9(2), 195–207. doi: 10.1006/nimg.1998.0396

Foucher, J. R., Otzenberger, H., & Gounot, D. (2003). The BOLD response and the gamma oscillations respond differently than evoked potentials: an interleaved EEG-fMRI study. BMC Neurosci, 4, 22. doi: 10.1186/1471-2202-4-22

Frauscher, B., von Ellenrieder, N., Dubeau, F., & Gotman, J. (2015). Scalp spindles are associated with widespread intracranial activity with unexpectedly low synchrony. Neuroimage, 105, 1–12. doi: 10.1016/j.neuroimage.2014.10.048

Gramfort, A., Papadopoulo, T., Olivi, E., & Clerc, M. (2010). OpenMEEG: opensource software for quasistatic bioelectromagnetics. Biomed Eng Online, 9, 45. doi: 10.1186/1475- 925X-9-45

Haegens, S., Barczak, A., Musacchia, G., Lipton, M. L., Mehta, A. D., Lakatos, P., & Schroeder, C. E. (2015). Laminar Profile and Physiology of the alpha Rhythm in Primary Visual, Auditory, and Somatosensory Regions of Neocortex. J Neurosci, 35(42), 14341–14352. doi: 10.1523/JNEUROSCI.0600-15.2015

Hagler, D. J., Jr. (2014). Optimization of retinotopy constrained source estimation constrained by prior. Human brain mapping, 35(5), 1815–1833. doi: 10.1002/hbm.22293

Hagler, D. J., Jr., & Dale, A. M. (2013). Improved method for retinotopy constrained source estimation of visual-evoked responses. Human brain mapping, 34(3), 665–683. doi: 10.1002/hbm.21461

Hagler, D. J., Jr., Halgren, E., Martinez, A., Huang, M., Hillyard, S. A., & Dale, A. M. (2009). Source estimates for MEG/EEG visual evoked responses constrained by multiple, retinotopically-mapped stimulus locations. Human brain mapping, 30(4), 1290–1309. doi: 10.1002/hbm.20597

Hämäläinen, M., Hari, R., Ilmoniemi, R. J., Knuutila, J., & Lounasmaa, O. V. (1993). Magnetoencephalography—theory, instrumentation, and applications to noninvasive studies of the working human brain. Reviews of modern Physics, 65(2), 413.

Hamalainen, M. S., & Ilmoniemi, R. J. (1994). Interpreting magnetic fields of the brain: minimum norm estimates. Med Biol Eng Comput, 32(1), 35–42. doi: 10.1007/bf02512476

Haufe, S., & Ewald, A. (2019). A Simulation Framework for Benchmarking EEG-Based Brain Connectivity Estimation Methodologies. Brain Topogr, 32(4), 625–642. doi: 10.1007/s10548-016-0498-y

Haxby, J. V., Gobbini, M. I., Furey, M. L., Ishai, A., Schouten, J. L., & Pietrini, P. (2001). Distributed and overlapping representations of faces and objects in ventral temporal cortex. Science, 293(5539), 2425–2430.

Heeger, D. J. (1992). Normalization of cell responses in cat striate cortex. Vis Neurosci, 9(2), 181–197. doi: 10.1017/s0952523800009640

Helmholtz, H. V. (1853). Ueber einige Gesetze der Vertheilung elektrischer Ströme in körperlichen Leitern, mit Anwendung auf die thierisch-elektrischen Versuche. Annalen der Physik, 165(7), 353–377.

Henrie, J. A., & Shapley, R. (2005). LFP power spectra in V1 cortex: the graded effect of stimulus contrast. J Neurophysiol, 94(1), 479–490. doi: 10.1152/jn.00919.2004

Henson, R. N., Mattout, J., Phillips, C., & Friston, K. J. (2009). Selecting forward models for MEG source-reconstruction using model-evidence. Neuroimage, 46(1), 168–176. doi: 10.1016/j.neuroimage.2009.01.062

Hermes, D., Miller, K. J., Vansteensel, M. J., Aarnoutse, E. J., Leijten, F. S., & Ramsey, N. F. (2012). Neurophysiologic correlates of fMRI in human motor cortex. Human brain mapping, 33(7), 1689–1699. doi: 10.1002/hbm.21314

Hermes, D., Miller, K. J., Wandell, B. A., & Winawer, J. (2015). Stimulus Dependence of Gamma Oscillations in Human Visual Cortex. Cereb Cortex, 25(9), 2951–2959. doi: 10.1093/cercor/bhu091

Hermes, D., Nguyen, M., & Winawer, J. (2017). Neuronal synchrony and the relation between the blood-oxygen-level dependent response and the local field potential. PLoS Biol, 15(7), e2001461. doi: 10.1371/journal.pbio.2001461

Hermes, D., Petridou, N., Kay, K. N., & Winawer, J. (2019). An image-computable model for the stimulus selectivity of gamma oscillations. Elife, 8. doi: 10.7554/eLife.47035

Huang, M. X., Mosher, J. C., & Leahy, R. M. (1999). A sensor-weighted overlapping-sphere head model and exhaustive head model comparison for MEG. Phys Med Biol, 44(2), 423–440. doi: 10.1088/0031-9155/44/2/010

Iivanainen, J., Stenroos, M., & Parkkonen, L. (2017). Measuring MEG closer to the brain: Performance of on-scalp sensor arrays. Neuroimage, 147, 542–553. doi: 10.1016/j.neuroimage.2016.12.048

Inverso, S. A., Goh, X. L., Henriksson, L., Vanni, S., & James, A. C. (2016). From evoked potentials to cortical currents: Resolving V1 and V2 components using retinotopy constrained source estimation without fMRI. Human brain mapping, 37(5), 1696–1709. doi: 10.1002/hbm.23128

Jas, M., Larson, E., Engemann, D. A., Leppakangas, J., Taulu, S., Hamalainen, M., & Gramfort, A. (2018). A Reproducible MEG/EEG Group Study With the MNE Software: Recommendations, Quality Assessments, and Good Practices. Front Neurosci, 12, 530. doi: 10.3389/fnins.2018.00530

Jeffreys, D. A. (1971). Cortical source locations of pattern-related visual evoked potentials recorded from the human scalp. Nature, 229(5285), 502–504. doi: 10.1038/229502a0

Jeffreys, D. A., & Axford, J. G. (1972a). Source locations of pattern-specific components of human visual evoked potentials. I. Component of striate cortical origin. Exp Brain Res, 16(1), 1–21.

Jeffreys, D. A., & Axford, J. G. (1972b). Source locations of pattern-specific components of human visual evoked potentials. II. Component of extrastriate cortical origin. Exp Brain Res, 16(1), 22–40.

Kay, K. N., Naselaris, T., Prenger, R. J., & Gallant, J. L. (2008). Identifying natural images from human brain activity. Nature, 452(7185), 352–355. doi: 10.1038/nature06713

Kay, K. N., Winawer, J., Mezer, A., & Wandell, B. A. (2013). Compressive spatial summation in human visual cortex. J Neurophysiol, 110(2), 481–494. doi: 10.1152/jn.00105.2013

Kay, K. N., Winawer, J., Rokem, A., Mezer, A., & Wandell, B. A. (2013). A two-stage cascade model of BOLD responses in human visual cortex. PLoS Comput Biol, 9(5), e1003079. doi: 10.1371/journal.pcbi.1003079

Kelly, S. P., Schroeder, C. E., & Lalor, E. C. (2013). What does polarity inversion of extrastriate activity tell us about striate contributions to the early VEP? A comment on Ales et al. (2010). Neuroimage, 76, 442–445. doi: 10.1016/j.neuroimage.2012.03.081

Kelly, S. P., Vanegas, M. I., Schroeder, C. E., & Lalor, E. C. (2013). The cruciform model of striate generation of the early VEP, re-illustrated, not revoked: a reply to Ales et al. (2013). Neuroimage, 82, 154–159. doi: 10.1016/j.neuroimage.2013.05.112

Keren, A. S., Yuval-Greenberg, S., & Deouell, L. Y. (2010). Saccadic spike potentials in gamma-band EEG: characterization, detection and suppression. Neuroimage, 49(3), 2248–2263. doi: 10.1016/j.neuroimage.2009.10.057

King, J. R., & Dehaene, S. (2014). Characterizing the dynamics of mental representations: the temporal generalization method. Trends Cogn Sci, 18(4), 203–210. doi: 10.1016/j.tics.2014.01.002

Krusienski, D. J., McFarland, D. J., & Wolpaw, J. R. (2012). Value of amplitude, phase, and coherence features for a sensorimotor rhythm-based brain-computer interface. Brain Res Bull, 87(1), 130–134. doi: 10.1016/j.brainresbull.2011.09.019

Kupers, E. R., Wang, H. X., Amano, K., Kay, K. N., Heeger, D. J., & Winawer, J. (2018). A non-invasive, quantitative study of broadband spectral responses in human visual cortex. PLoS One, 13(3), e0193107. doi: 10.1371/journal.pone.0193107

Kybic, J., Clerc, M., Abboud, T., Faugeras, O., Keriven, R., & Papadopoulo, T. (2005). A common formalism for the integral formulations of the forward EEG problem. IEEE Trans Med Imaging, 24(1), 12–28.

Liu, T., Heeger, D. J., & Carrasco, M. (2006). Neural correlates of the visual vertical meridian asymmetry. J Vis, 6(11), 1294–1306. doi: 10.1167/6.11.12

Lopes, M. A., Junges, L., Tait, L., Terry, J. R., Abela, E., Richardson, M. P., & Goodfellow, M. (2020). Computational modelling in source space from scalp EEG to inform presurgical evaluation of epilepsy. Clin Neurophysiol, 131(1), 225–234. doi: 10.1016/j.clinph.2019.10.027

Manning, J. R., Jacobs, J., Fried, I., & Kahana, M. J. (2009). Broadband shifts in local field potential power spectra are correlated with single-neuron spiking in humans. J Neurosci, 29(43), 13613–13620. doi: 10.1523/JNEUROSCI.2041-09.2009

Mante, V., Bonin, V., & Carandini, M. (2008). Functional mechanisms shaping lateral geniculate responses to artificial and natural stimuli. Neuron, 58(4), 625–638. doi: 10.1016/j.neuron.2008.03.011

Miller, K. J., Honey, C. J., Hermes, D., Rao, R. P., denNijs, M., & Ojemann, J. G. (2014). Broadband changes in the cortical surface potential track activation of functionally diverse neuronal populations. Neuroimage, 85 *Pt* *2*, 711–720. doi: 10.1016/j.neuroimage.2013.08.070

Miller, K. J., Leuthardt, E. C., Schalk, G., Rao, R. P., Anderson, N. R., Moran, D. W., … Ojemann, J. G. (2007). Spectral changes in cortical surface potentials during motor movement. J Neurosci, 27(9), 2424–2432. doi: 10.1523/JNEUROSCI.3886-06.2007

Miller, K. J., Sorensen, L. B., Ojemann, J. G., & den Nijs, M. (2009). Power-law scaling in the brain surface electric potential. PLoS Comput Biol, 5(12), e1000609. doi: 10.1371/journal.pcbi.1000609

Milstein, J., Mormann, F., Fried, I., & Koch, C. (2009). Neuronal shot noise and Brownian 1/f2 behavior in the local field potential. PLoS One, 4(2), e4338. doi: 10.1371/journal.pone.0004338

Moradi, F., Liu, L. C., Cheng, K., Waggoner, R. A., Tanaka, K., & Ioannides, A. A. (2003). Consistent and precise localization of brain activity in human primary visual cortex by MEG and fMRI. Neuroimage, 18(3), 595–609. doi: 10.1016/s1053-8119(02)00053-8

Movshon, J. A., Thompson, I. D., & Tolhurst, D. J. (1978). Receptive field organization of complex cells in the cat’s striate cortex. J Physiol, 283, 79–99. doi: 10.1113/jphysiol.1978.sp012489

Mukamel, R., Gelbard, H., Arieli, A., Hasson, U., Fried, I., & Malach, R. (2005). Coupling between neuronal firing, field potentials, and FMRI in human auditory cortex. Science, 309(5736), 951–954. doi: 10.1126/science.1110913

Musall, S., von Pfostl, V., Rauch, A., Logothetis, N. K., & Whittingstall, K. (2014). Effects of neural synchrony on surface EEG. Cereb Cortex, 24(4), 1045–1053. doi: 10.1093/cercor/bhs389

Muthukumaraswamy, S. D. (2013). High-frequency brain activity and muscle artifacts in MEG/EEG: a review and recommendations. Front Hum Neurosci, 7, 138. doi: 10.3389/fnhum.2013.00138

Naselaris, T., Kay, K. N., Nishimoto, S., & Gallant, J. L. (2011). Encoding and decoding in fMRI. Neuroimage, 56(2), 400–410. doi: 10.1016/j.neuroimage.2010.07.073

Nasiotis, K., Clavagnier, S., Baillet, S., & Pack, C. C. (2017). High-resolution retinotopic maps estimated with magnetoencephalography. Neuroimage, 145(Pt A), 107–117. doi: 10.1016/j.neuroimage.2016.10.017

Neymotin, S. A., Daniels, D. S., Caldwell, B., McDougal, R. A., Carnevale, N. T., Jas, M., … Jones, S. R. (2020). Human Neocortical Neurosolver (HNN), a new software tool for interpreting the cellular and network origin of human MEG/EEG data. Elife, 9. doi: 10.7554/eLife.51214

Norcia, A. M., Appelbaum, L. G., Ales, J. M., Cottereau, B. R., & Rossion, B. (2015). The steady-state visual evoked potential in vision research: A review. J Vis, 15(6), 4. doi: 10.1167/15.6.4

Norcia, A. M., & Tyler, C. W. (1985). Spatial frequency sweep VEP: visual acuity during the first year of life. Vision Res, 25(10), 1399–1408. doi: 10.1016/0042-6989(85)90217- 2

Oostenveld, R., Fries, P., Maris, E., & Schoffelen, J. M. (2011). FieldTrip: Open source software for advanced analysis of MEG, EEG, and invasive electrophysiological data. Comput Intell Neurosci, 2011, 156869. doi: 10.1155/2011/156869

Pantazis, D., Fang, M., Qin, S., Mohsenzadeh, Y., Li, Q., & Cichy, R. M. (2018). Decoding the orientation of contrast edges from MEG evoked and induced responses. Neuroimage, 180(Pt A), 267–279. doi: 10.1016/j.neuroimage.2017.07.022

Pfurtscheller, G., & Cooper, R. (1975). Frequency dependence of the transmission of the EEG from cortex to scalp. Electroencephalogr Clin Neurophysiol, 38(1), 93–96.

Poghosyan, V., & Ioannides, A. A. (2007). Precise mapping of early visual responses in space and time. Neuroimage, 35(2), 759–770. doi: 10.1016/j.neuroimage.2006.11.052

Ray, S., & Maunsell, J. H. (2011). Different origins of gamma rhythm and high-gamma activity in macaque visual cortex. PLoS Biol, 9(4), e1000610. doi: 10.1371/journal.pbio.1000610

Regan, D. (1989). Human brain electrophysiology. Evoked potentials and evoked magnetic fields in science and medicine.

Riera, J. J., Ogawa, T., Goto, T., Sumiyoshi, A., Nonaka, H., Evans, A., . . . Kawashima, R. (2012). Pitfalls in the dipolar model for the neocortical EEG sources. J Neurophysiol, 108(4), 956–975. doi: 10.1152/jn.00098.2011

Robinson, A. K., Venkatesh, P., Boring, M. J., Tarr, M. J., Grover, P., & Behrmann, M. (2017). Very high density EEG elucidates spatiotemporal aspects of early visual processing. Sci Rep, 7(1), 16248. doi: 10.1038/s41598-017-16377-3

Rust, N. C., Schwartz, O., Movshon, J. A., & Simoncelli, E. P. (2005). Spatiotemporal elements of macaque v1 receptive fields. Neuron, 46(6), 945–956. doi: 10.1016/j.neuron.2005.05.021

Sharon, D., Hamalainen, M. S., Tootell, R. B., Halgren, E., & Belliveau, J. W. (2007). The advantage of combining MEG and EEG: comparison to fMRI in focally stimulated visual cortex. Neuroimage, 36(4), 1225–1235. doi: 10.1016/j.neuroimage.2007.03.066

Simoncelli, E. P., & Heeger, D. J. (1998). A model of neuronal responses in visual area MT. Vision Res, 38(5), 743–761. doi: 10.1016/s0042-6989(97)00183-1

Stenroos, M., Hunold, A., & Haueisen, J. (2014). Comparison of three-shell and simplified volume conductor models in magnetoencephalography. Neuroimage, 94, 337–348. doi: 10.1016/j.neuroimage.2014.01.006

Stenroos, M., & Nummenmaa, A. (2016). Incorporating and Compensating Cerebrospinal Fluid in Surface-Based Forward Models of Magneto- and Electroencephalography. PLoS One, 11(7), e0159595. doi: 10.1371/journal.pone.0159595

Steriade, M., McCormick, D. A., & Sejnowski, T. J. (1993). Thalamocortical oscillations in the sleeping and aroused brain. Science, 262(5134), 679–685. doi: 10.1126/science.8235588

Tadel, F., Baillet, S., Mosher, J. C., Pantazis, D., & Leahy, R. M. (2011). Brainstorm: a user-friendly application for MEG/EEG analysis. Comput Intell Neurosci, 2011, 879716. doi: 10.1155/2011/879716

Tallon-Baudry, C., & Bertrand, O. (1999). Oscillatory gamma activity in humans and its role in object representation. Trends Cogn Sci, 3(4), 151–162. doi: 10.1016/s1364-6613(99)01299-1

Uhlhaas, P. J., & Singer, W. (2006). Neural synchrony in brain disorders: relevance for cognitive dysfunctions and pathophysiology. Neuron, 52(1), 155–168. doi: 10.1016/j.neuron.2006.09.020

Van Der Tweel, L. H., & Lunel, H. F. (1965). Human Visual Responses to Sinusoidally Modulated Light. Electroencephalogr Clin Neurophysiol, 18, 587–598. doi: 10.1016/0013-4694(65)90076-3

Wang, L., Mruczek, R. E., Arcaro, M. J., & Kastner, S. (2015). Probabilistic Maps of Visual Topography in Human Cortex. Cereb Cortex, 25(10), 3911–3931. doi: 10.1093/cercor/bhu277

Winawer, J., Kay, K. N., Foster, B. L., Rauschecker, A. M., Parvizi, J., & Wandell, B. A. (2013). Asynchronous broadband signals are the principal source of the BOLD response in human visual cortex. Curr Biol, 23(13), 1145–1153. doi: 10.1016/j.cub.2013.05.001

Yuval-Greenberg, S., & Deouell, L. Y. (2009). The broadband-transient induced gamma-band response in scalp EEG reflects the execution of saccades. Brain Topogr, 22(1), 3–6. doi: 10.1007/s10548-009-0077-6

Yuval-Greenberg, S., Tomer, O., Keren, A. S., Nelken, I., & Deouell, L. Y. (2008). Transient induced gamma-band response in EEG as a manifestation of miniature saccades. Neuron, 58(3), 429–441. doi: 10.1016/j.neuron.2008.03.027

